# Olive phenolic compounds, potent tankyrase 1 inhibitor exhibit anti colon cancer effects by blocking Wnt/β-catenin pathway

**DOI:** 10.1101/2024.05.08.593091

**Authors:** Arindam Sain, Soumen Barman, Dipsikha Khamrai, M Srinivas, Prerna Singh, Bhargav Simpi, Mayuri Varshney, Debdut Naskar

## Abstract

Colon cancer (CC) needs special attention to develop novel targets and therapeutic approaches, as the total number of morbidity and mortality is growing rapidly. Although Olive has been long reported for its anti-CC activities but the mode is not completely understood yet. Here, we targeted tankysare 1 (*TNKS*), an important mediator of the Wnt/β-catenin pathway, crucial in colon carcinogenesis. Bioinformatics analysis revealed that the mutation and copy number alterations of *TNKS* is found to be 12% whereas the proteins expression of *TNKS* is moderate to high in tumor groups vs normal in CC datasets. On the more, *TNKS* and its positively associated genes are found to be concomitant with several carcinogenesis related biological processes and signaling pathways. Further, three olive phytochemicals (apigenin, luteolin, and quercetin), which were shortlisted by employing filters like drug likeness and toxicity screening, molecular docking and finally molecular dynamics simulations, can target tankyrase 1. These ployphenols demonstrated cytotoxic effect in CC (HT-29) cells either alone or in combination. Further, these compounds successfully reduced β-catenin level and its target genes (*CCND1, CDK4*, and *MYC*). Overall, inhibition of the Wnt/β-catenin pathway by targeting tankyrase 1 was revealed as one of the anti-CC mechanisms of olive polyphenols.

## 1. Introduction

Colon cancer (CC) casts a looming shadow on global healthcare, ranking third in number of cases of diagnoses worldwide and second in cancer-related deaths. Its impact extends beyond western countries, affecting populations globally (1). According to the GLOBOCAN database, only in 2020, the estimated occurrence of colon cancer was 1.9 million, with over 0.9 million related deaths (2). Apart from the genetic predisposition (e.g., mutation), possible reasons behind the statistics include changes in lifestyle like fewer physical activities, low-fiber and high-fat food intake, smoking, and alcoholism (3). Unfortunately, the number of effective therapeutic targets and modalities is very limited. Conventional therapies, such as surgery and chemoradiotherapy, are frequently afflicted with serious side effects (4). In contrast, targeted therapies, such as anti-EGFR or anti-VEGF drugs, are typically unfeasible for the majority of patients due to their high expenses (5). Therefore, the search for new targets in colon cancer is obvious, and different signaling pathways have been targeted recently; for example, cetuximab or panitumumab are being used in CC, which targets the epidermal growth factor receptor (EGFR) (6).

The disruption of cellular signaling pathways is responsible for numerous disorders, including CC. Signaling pathways such as EGFR/MAPK, TGF-β, Notch, PI3K/AKT, and Wnt/β-catenin are crucial in the development of CC (7). In the progression of CC, the Wnt/β-catenin pathway assumes a crucial role, influencing processes such as cell proliferation, differentiation, survival, and apoptosis (8). Different components of this pathway have been targeted to regulate or modify the Wnt/β-catenin pathway in CC. Tankyrases (1 and 2) receive major attention as a target to modulate the Wnt/β-catenin signaling due to their ability to play an essential role in mediating the destruction of the AXINs (AXIN1 and AXIN2), which negatively regulate the oncogenic Wnt/β-catenin signaling (9).

PAR (Poly ADP-Ribose)-dependent ubiquitinylation (PARdU) of AXIN by tankyrase renders the AXIN degradation by the proteasome (10). It helps accumulation of β-catenin in the cytoplasm and subsequent translocation in the nucleus, which causes the progression of proto-oncogenic signaling. Therefore, inhibition or modulation of the tankyrase activity can be an attractive target to control excessive Wnt/β-catenin pathway activity and prevent CC initiation or progression.

The Olive tree (*Olea europaea*) is known for its medicinal properties due to the presence of a diverse range of active compounds, mainly phenolic compounds. The Mediterranean diet, primarily based on olive oil, is known for its preventive role against CC. Olive oil phenolic compounds were previously shown to exert anti-cancerous activities in different cancers (11–13). However, a complete picture of the interaction of cellular signaling pathways with phenolic compounds still needs to be provided. This study aimed to divulge the precise anti-cancer mechanism of olive phenolic compounds by studying the interaction between the tankyrase 1/Wnt/β-catenin pathway and olive compounds to get more insight into olive’s anti-CC roles and also identify a potent tankyrase-1 inhibitor from olive-derived compounds.

## 2. Materials and methods

### 2.1. Genetic Alteration analysis of the TNKS

The genetic alteration and the mutation frequency of the *TNKS* (translates Tankyrase-1) in colon cancer within the TCGA database were analyzed and explored using the CBioportal database (https://www.cbioportal.org) (14). CBioportal contains vast mutation and genetic alterations data across different datasets of patients and cancer types across different countries, thereby providing a multidimensional insight into the cancer. Different colon cancer datasets were referred for the study of genetic anomalies and the copy number alterations of *TNKS* to further validate its significance and the major role played by *TNKS* in colon carcinogenesis. *TNKS* expression was further checked through the Human Protein Atlas (https://www.proteinatlas.org/).

### 2.2. Genetic association analysis of TNKS and functional enrichment analysis

The LinkedOmics web portal (15) was used to reveal the top 50 significantly associated genes of *TNKS* (both positively and negatively correlated genes) by applying the Spearman correlation test. Further, the LinkInterpreter module of the LinkedOmics suite was executed to get an idea of the relevant biological processes, molecular functions, and KEGG and Reactome pathway enrichment with the top 50 positively associated genes of *TNKS*.

### 2.3. Data Curation and Drug Screening

A total of 118 compounds were selected from the *Olea europaea* plant after a literature review, and their presence in *O. europaea* was further rechecked via the Olivamine (OliveNet) database (16). For this study, only phenolic compounds were taken into consideration. The phenolic compounds obtained were subjected to a drug-likeness screening by utilizing the Molsoft Drug Likeness (https://molsoft.com/mprop) software and applying the Lipinskis ‘rule of 5’ filters (Hydrophobicity, Drug Likeness, Molecular Weight, MolLogP, MolPSA), by the Molinspiration Cheminformatics Server (https://www.molinspiration.com/cgi-bin/proper). The Canonical SMILES structure of each of the compounds was obtained from PubChem (https://pubchem.ncbi.nlm.nih.gov) and the HMDB database (17). Compounds that violate Lipinski’s rule (more than one violation) were removed from further analysis.

### 2.4. Toxicity studies

Toxicity prediction or studies are a crucial part of the drug development process. The toxicity profiling of the screened polyphenols was done using the pkCSM server (https://biosig.lab.uq.edu.au/pkcsm/). Mainly six parameters (carcinogenicity, AMES toxicity, hERG-I or hERG-II inhibitor, hepatotoxicity, and skin sensitization) were considered for assessing the toxicity of a particular polyphenol. The compounds that passed all of the above-mentioned parameters were selected for the next stage of drug screening.

### 2.5. Protein target preparation

The RCSB PDB database (https://www.rcsb.org/) was inspected to find a suitable 3D structure for our target protein, tankyrase 1. The PDB ID: 3UDD [1.95 Å] was found to be the most suitable one, having the highest resolution and encompassing the catalytic site. The 3D crystal structure was cleared of any water molecule, ligand group, or any other kind of heteroatom present. Polar hydrogen and gasteiger charges were added to stabilize the protein crystal structure and make it ready for molecular docking (18).

### 2.6. Ligand preparation

The three-dimensional conformation of each of the phenolic compounds was obtained from the PubChem database in the SDF format. The SDF format was converted to the PDB format using OpenBabel 2.4.1. (19). The PDB formats were then processed and converted into the pdbqt format using the AutoDock Tools (v4.2.) for Vina docking (20). Energy minimization of the ligand molecule was done with the Avogadro 1.2.0 software (https://avogadro.cc/).

### 2.7. Exploring protein-ligand interaction by molecular Docking and Visualization

The molecular docking was executed using the AutoDock Vina software (21), and the grid box was set in such a way that it would cover the entire active site responsible for the catalytic activity of the enzyme, so that the ligand molecule is targeted to bind with the active site of the enzyme molecule. The docking conformations with the lowest binding free energy were selected for the final docking results. Finally, the results were visualized and demonstrated as a 2D representation through the BIOVIA Discovery Studio Visualizer v21.1.0.20298.

### 2.8. Molecular Dynamics Simulation

100 ns of MD simulations were executed via the GROMACS (version 2018.3) package by using the GROMOS96 54A7 force field for the top-ranked docked conformations of the molecular docking (22). MD simulations of the protein-peptide complexes were set up after creating the topology parameters for both proteins and ligands through GROMACS and PRODRG2 servers, respectively (23). The solvation of the prepared protein-ligand complexes was executed following standard protocol. MD simulations of the ligand-tankyrase 1 were carried out by following the parameters: 300K temperature, 1 atm pressure, and a 2 fs time step for 100 ns. The calculation of the RMSD (root mean square deviation), RMSF (root mean square fluctuation), and Rg (radius of gyration) values was done by analyzing the final MD trajectories with the help of standard GROMACS functions.

### 2.9. Binding free energy calculations (MM-PBSA method)

The g_mmpbsa tool was used to determine the binding free energy of protein-ligand complexes derived from MD trajectories. Following simulation, the MM-PBSA (Molecular Mechanics Poisson Boltzmann Surface Area) module was run to calculate the binding free energy, which estimates Gibb’s free energy of binding using the previously given equation. (24).

### 2.10. Cell culture and Cytotoxicity assay

The HT-29 colon cancer cell line was purchased from the National Centre for Cell Science (NCCS), Pune, India, and cultured in DMEM media with 10% fetal bovine serum and 1X Gibco™ Antibiotic-Antimycotic (Cat No. 15240062) within a humidified chamber (5% CO2 and 37°C). Apigenin, luteolin, and quercetin (all from Sigma-Aldrich) were dissolved in DMSO (dimethyl sulfoxide, HIMEDIA, India). The top-selected compounds (apigenin, luteolin, and quercetin) from the *in-silico* analysis was tested in cell culture to reveal the potential cytotoxic effects of olive compounds against CC cells (HT-29). We performed the MTT assay with different doses of apigenin, luteolin, and quercetin and also applied these compounds in combination to check the cytotoxicity of these compounds following the standard protocol previously described (25).

### 2.11. Colony formation assay

HT-29 cells were seeded in a 12-well plate at a density of 1000 cells per well. Cells were treated with apigenin, luteolin, quercetin, and a combination of these three compounds for 48 hours. Then cells were grown for another 2 weeks until visible colonies were grown (a colony should contain at least 50 cells). The plate was then fixed with a solution of methanol and glacial acetic acid in a 3:1 ratio, and finally stained with a crystal violet solution (0.5%) (26).

### 2.12. Acridine orange/ethidium bromide (AO/EtBr) double staining

HT-29 cells were seeded in a 24-well plate at a density of 50000 cells/well, and cells were treated with apigenin (80 μM), quercetin (80 μM), or luteolin (70 μM) for a duration of 48 h. Cells were then washed with 1X PBS and added working solution of AO/EtBr (prepared by mixing 1μL of AO (5 mg/mL) and 1μL EtBr (3 mg/mL) in 1mL of 1x PBS) and immediately checked under a fluorescence microscope (27).

### 2.13. Quantitative real-time PCR analysis to check mRNA expression

The mRNA expression of relevant genes of the Wnt/β-catenin pathway in HT-29 cells was checked after treatment with apigenin, luteolin, and quercetin alone or in combinations. RNA was extracted from cells using TRIzol reagent (Invitrogen) once the treatment period was over, followed by cDNA synthesis using a cDNA synthesis kit (BioBharati LifeScience, India), following the manufacturer’s instructions. SYBR green PCR master mix (Applied Biosystem) and respective primer pairs (*ACTB* - F: 5’-GGATTCCTATGTGGGCGACGAG-3’, R: 5’-GAGCCACACGCAGCTCATTG-3’; *CCND1* - F: 5’ - GAGGGACGCTTTGTCTGTCG – 3’, R: 5’ – ACATGTTGGTGCTGGGAAGC – 3’; *CDK4* – F: 5’ – AACTCTGAAGCCGACCAGTTG – 3’, R: 5’-AGTCAGCATTTCCAGCAGCAG – 3’; *MYC* – F: 5’ – GAAGGAATGGCAGAAGGCAG – 3’, R: 5’ – AGAATTCCTGGGTTTGGAGTG – 3’) of target genes were utilized to quantify mRNA expression with an StepOneTM real-time PCR (Applied Biosystem) machine. The *ACTB* (β-Actin) expression was used to normalize the expression. Relative expression of target genes was quantified by the 2^−ΔΔCT^ method based on C_T_ values (28).

### 2.14. Immunoblot analysis

Protein level expression of target proteins was checked in HT-29 cells after treatment with the respective concentrations of the interventions and their combinations or vehicle control. Cells were harvested in ice-cold lysis buffer (50 mM Tris-HCL, pH 8.0, 150 mM NaCl, 1% Triton X100, 1 mM EDTA, 0.1% SDS, and 20 mM PMSF, 5 mM NaF, 1 mM Na_3_VO_4_, and 0.5% sodium deoxycholate). Bradford reagent (Sigma-Aldrich) was used to quantify protein from the cell lysate by using a multi-plate reader (Thermo Scientific). Equal amount of proteins from each sample was resolved by SDS-PAGE and further probed with specific antibodies against β-catenin (Invitrogen). Antibody against β-actin, the housekeeping gene was employed as internal control in immunoblot analysis. After primary antibody incubation, anti-mouse secondary antibody (Cell Signaling Technology) was used to probe target proteins with the BCIP®/NBT Liquid Substrate (Sigma-Aldrich). ImageJ software (National Institutes of Health) was employed for densitometry analysis to quantify target proteins (29).

### 2.15. Generation of CC spheroids and treatment with selected olive Phytochemicals

CC spheroids were generated using HT-29 cells grown in ultra-low attachment Multidishes (Nunc™). For spheroid culture specific serum free DMEM-F12 media (SFM) was used, supplemented with FGF, fibroblast growth factor (10 ng/mL, Merck), EGF, epidermal growth factor (20 ng/mL, Gibco™), B-27 supplements (at 1X concentration, Gibco™) and N-2 supplement (1X concentration, Gibco™, 1X). Cells were seeded at a concentration of 5000 cells/well. After 12 h of overnight incubation, cells which were started forming spheroid by that time were treated with appropriate phytochemicals and/or their combinations (30).

### 2.16. Statistical analysis

GraphPad Prism 6 software was utilized to analyze results by paired Student t test. Results were expressed as mean of ± SEM; Significance was annotated by p value, *p ≤ 0.05, **p ≤ 0.01, *** p ≤ 0.001.

## 3. Results and discussion

### 3.1. Genetic alteration, expression, and association study of TNKS

Mutation and structural variation analysis of the *TNKS* on the CBioportal server revealed a high alteration frequency of the gene within different datasets. We analyzed four different datasets: Colon Cancer (CPTAC-2 Prospective, Cell 2019), Colon Cancer (Sidra-LUMC AC-ICAM, Nat Med 2023), Colorectal Adenocarcinoma (TCGA, Pan Cancer Atlas), and Colorectal Adenocarcinoma (TCGA, Firehose Legacy). 12% alteration frequency was observed within the CPTAC-2 Prospective (Cell 2019) dataset (110 patients) which includes missense mutations, truncating mutations, and deep deletions. Analysis of the Colorectal Adenocarcinoma (378 patients) (TCGA, Pan Cancer Atlas) dataset revealed a 10% alteration frequency of the *TNKS*. The other two datasets, Colon Cancer (Sidra-LUMC AC-ICAM, Nat Med 2023, 348 patients) and Colorectal Adenocarcinoma (TCGA, Firehose Legacy, 392 patients) having *TNKS* alterations frequency of 9% and 8%, respectively (Figure 1). Any correlation between *TNKS* alteration frequency and patients’ sex was also considered in all these four datasets. Interestingly, we found males were affected (62.04%) more than females (37.96%) when calculated using all four datasets (Supplementary Figure 1). From the HPA server, it was revealed that tankyrase 1 protein expression level in CRC was moderate to high in CRC patients (∼ 64% of patients) and CRC only comes third after liver and ovarian cancer among all cancers (Figure 2A). The normalized gene expression level (nTPM) was also checked in different CRC cell lines, and an expression level of nTPM 3.9 to 16.4 was observed in different CRC cell lines (Figure 2B) which implies overall good expression value of TNKS in different cell lines and its crucial role in CRC cell line models. With the immunohistochemistry images available from the HPA server, it was revealed that tankyrase 1 staining was high in tumor tissue compared to normal tissue (Figure 2C).

**Figure 1:**
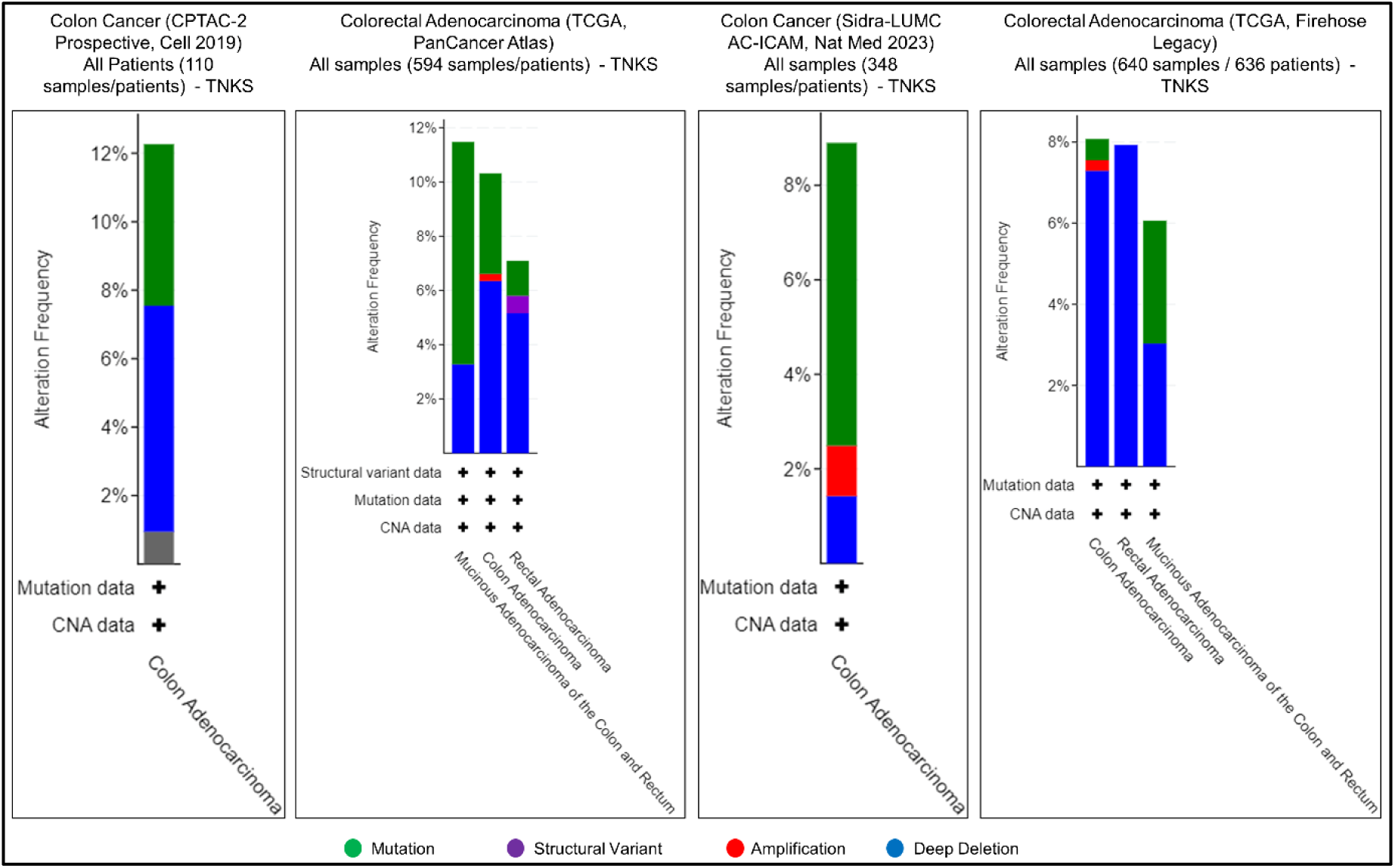
Genomic alterations (mutations and copy number alterations) of *TNKS* from different colon cancer databases. Alterations (mutations and copy number alterations) frequency of 12%, 10%, 9%, and 8% were observed from four different datasets.

**Figure 2:**
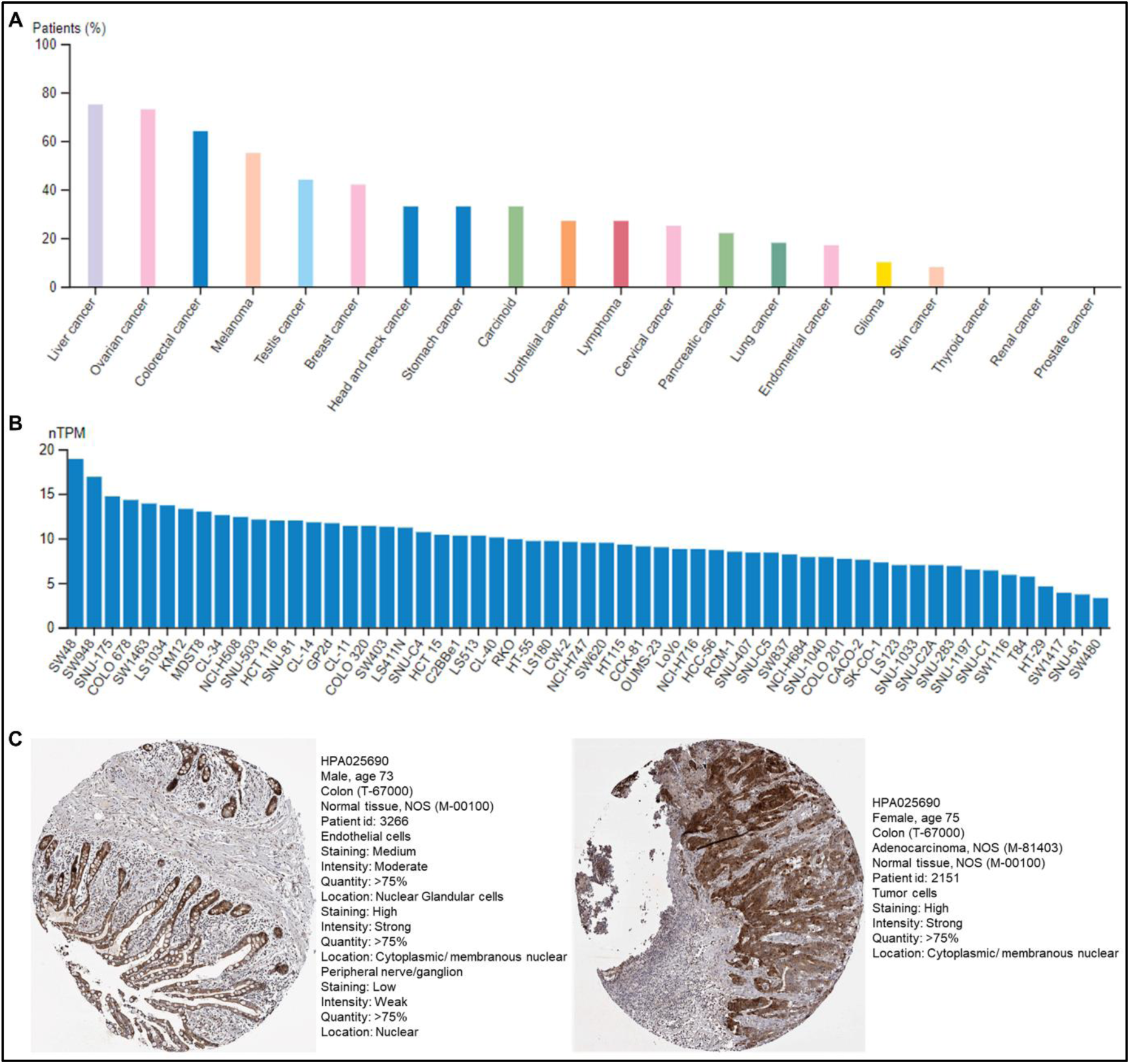
Expression analysis of tankyrase 1 from the Human Protein Atlas database. A: Expression of tankyrase 1 in different cancers, B: expression level of tankyrase 1 in different CRC cell lines, C: representative immunohistochemistry images of tankyrase 1 protein expression in normal (left side) and tumour tissue (right side) sections.

From the genetic association study executed on the LinkedOmics site (LinkFinder module), the top 50 significantly associated genes (both positive and negative) of TNKS were revealed (Figures 3A, 3B, and 3C). The positively associated genes were further subjected to functional enrichment analyses by the LinkInterpreter module, and these genes were revealed to control several important biological processes (regulation of immune system processes, cell development) and molecular functions (transcription regulator activity, kinase binding) linked to carcinogenesis. Further, these genes were enriched in significant KEGG and Reactome pathways, including the cGMP-PKG signaling pathway, the PI3K-Akt signaling pathway, the MAPK signaling pathway, extracellular matrix organization, and signaling by receptor tyrosine kinases (Figure 4A, 4B, 4C, and 4D). Altogether, the high alteration frequency and higher protein expression of tankyrase 1 make it an interesting target to overcome the burden of colon cancer. Moreover, tankyrase 1 and its positively associated genes are related to several crucial biological processes and signaling pathways; alteration of those might lead to carcinogenesis, which indicates the crucial role of tankyrase 1 in CC.

**Figure 3:**
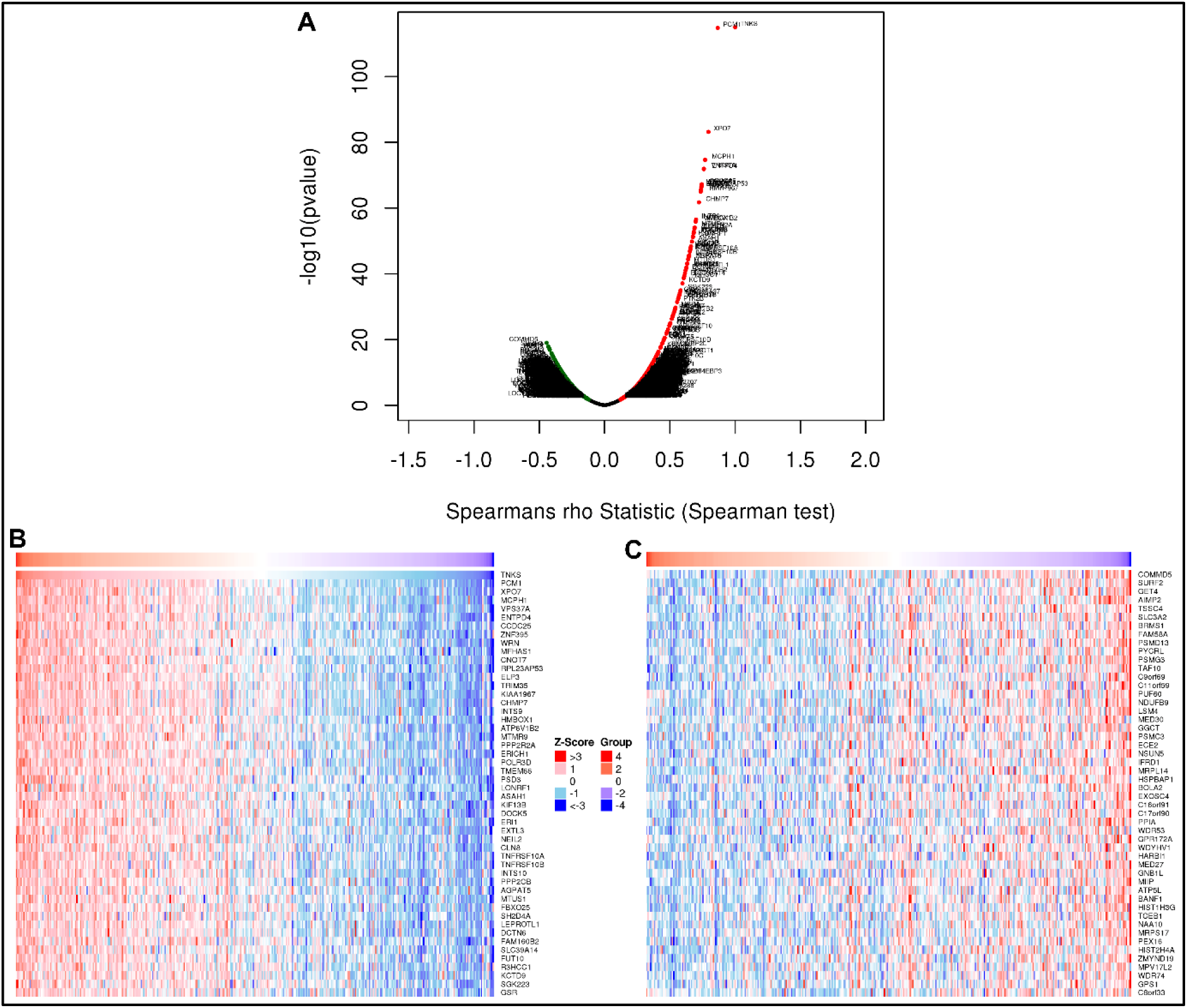
Genetic association study of *TNKS* (tankyrase 1) from the LinkedOmics database. A: *TNKS* association results (volcano graph). B: top 50 positively correlated significant genes of *TNKS* (heat map). C: top 50 negatively correlated genes of *TNKS* (heat map).

**Figure 4:**
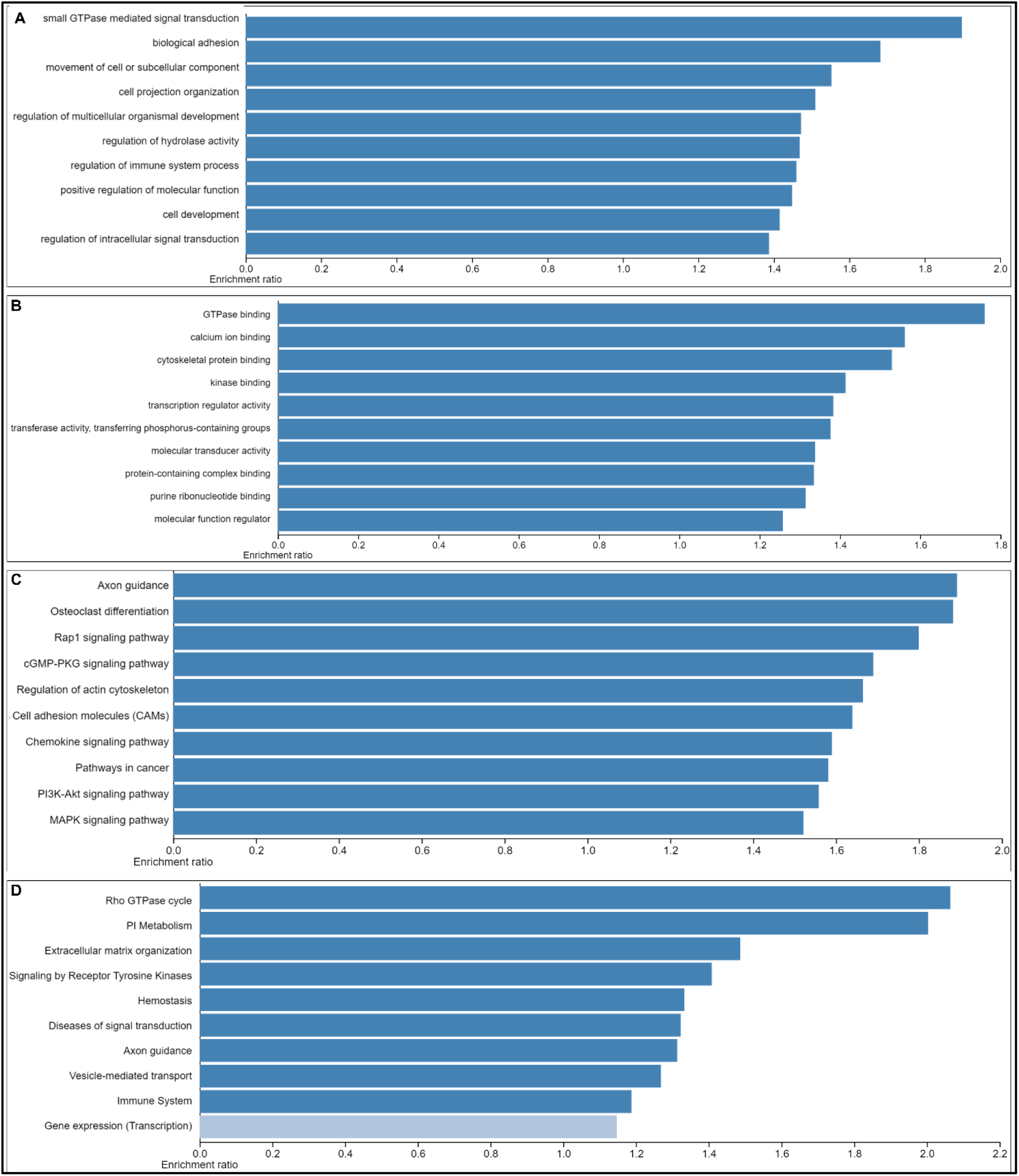
Functional enrichment analysis of the top 50 positively associated genes of TNKS. Biological processes (A), molecular functions (B), KEGG pathways (C), and Reactome pathways (D).

### 3.2. Drug likeness screening facilitated the identification of suitable drug-like phenols present in olive

In order to search for potential drug candidates which can target tankyrase in CC, we focused on olive and its phytoconstitutes as it is well-known for its anti-colon cancer activities but overall mechanism is yet to be fully understood. To reveal the possible anti-colon cancer mechanism of olive, a total of 118 compounds were curated from *O. europaea* (Supplementary table 1). After screening the compounds with the drug-likeness filters, a set of 18 compounds were selected for further analysis. The drug likeness score of 0.30 was considered the cut-off score for the selection of compounds (Table 1).

**Table 1:** List of shortlisted compounds from Olive after Drug Likeness analysis.

| Sl. No. | Compound Name | PubChem CID | Drug Likeness Score |
| --- | --- | --- | --- |
| 1. | Apigenin | 5280443 | 0.39 |
| 2. | Apigenin 7-O glucoside | 12304093 | 0.59 |
| 3. | Chrysoeriol | 5280666 | 0.30 |
| 4. | Berchemol | 14521044 | 0.65 |
| 5. | Gentisic Acid | 3469 | 0.30 |
| 6. | 3,4-Dihydroxyphenylglycol | 91528 | 0.71 |
| 7. | Chlorogenic Acid | 1794427 | 0.79 |
| 8. | Cornoside | 3084796 | 0.61 |
| 9. | Eriodictyol | 90657147 | 0.91 |
| 10. | Hesperetin | 72281 | 0.59 |
| 11. | Hydroxytyrosil elenolate | 101109285 | 0.49 |
| 12. | Loganic Acid | 89640 | 1.13 |
| 13. | Loganin | 87691 | 0.88 |
| 14. | Luteolin | 5280445 | 0.38 |
| 15. | 6-Methoxyluteolin | 5317284 | 0.44 |
| 16. | Quercetin | 5280343 | 0.52 |
| 17. | Taxifolin | 12309272 | 1.00 |
| 18. | Rosmarinic Acid | 5281792 | 0.37 |

### 3.3. Toxicity analysis revealed potentially toxic phenols

Toxicity profiling of the phytochemicals was done by employing the pkCSM server (https://biosig.lab.uq.edu.au/pkcsm/) to check AMES toxicity, hERG-I, hHERG-II, hepatotoxicity, and skin sensitization. This helped to uncover any potential toxicity of the concerned compounds. Seventeen compounds were qualified as potential drug candidates without any toxicity, and toxic compound hydroxytyrosil elenolate was subsequently excluded from downstream analysis (Table 2). Therefore, among all the phenolic compounds these compounds might be responsible behind the protective action of olive against carcinogenesis with favorable drug likeness profile and without any associated toxicity.

**Table 2:** List of compounds selected after the toxicity filtering. Compounds possess toxicity highlighted in italic and bold.

| Sl. No. | Compound Name | AMES toxicity | hERG-I inhibitor | hERG-II inhibitor | Hepato toxicity | Skin sensitization |
| --- | --- | --- | --- | --- | --- | --- |
| 1. | Apigenin | - | - | - | - | - |
| 2. | Berchemol | - | - | - | - | - |
| 3. | Apigenin 7-O glucoside | - | - | - | - | - |
| 4. | Chlorogenic Acid | - | - | - | - | - |
| 5. | 3,4-Dihydroxyphenylglycol | - | - | - | - | - |
| 6. | Eridictyol | - | - | - | - | - |
| 7. | Hesperitin | - | - | - | - | - |
| 8. | Loganic Acid | - | - | - | - | - |
| 9. | ***Hydroxytyrosil elenolate*** | + | - | - | - | - |
| 10. | Loganin | - | - | - | - | - |
| 11. | Luteolin | - | - | - | - | - |
| 12. | Quercetin | - | - | - | - | - |
| 13. | Taxifolin | - | - | - | - | - |
| 14. | Gentisic Acid | - | - | - | - | - |
| 15. | Cornoside | - | - | - | - | - |
| 16. | 6-Methoxyluteolin | - | - | - | - | - |
| 17. | Rosmarinic Acid | - | - | - | - | - |
| 18. | Chrysoeriol | - | - | - | - | - |

### 3.4. Molecular Docking analysis revealed strong protein-ligand interactions between Tankyrase and olive phytochemicals

Next, we performed molecular docking of the target enzyme tankyrase 1 with the selected olive phenols (qualified both drug likeness and toxicity filter) to reveal protein-ligand interactions, and XAV939, a known tankyrase inhibitor (31), was employed as a reference compound in our study. Majority of the tested compounds showed better binding affinity than the reference compound (Supplementary table 2). Among all screened compounds, the top five ranked compounds were selected based on their docking scores (binding affinity). The docking score varied from -9.4 Kcal/mol to -10.0 Kcal/mol (Table 3a). Eriodyctiol showed the best docking score with tankyrase 1, followed by luteolin, apigenin, hesperitin, 6-methoxyluteolin, and quercetin. These polyphenols were found to interact with the PARP catalytic site of the tankyrase1 enzyme. The primary interactions found were hydrogen bonds and non-covalent interactions (Figure 5).

**Figure 5:**
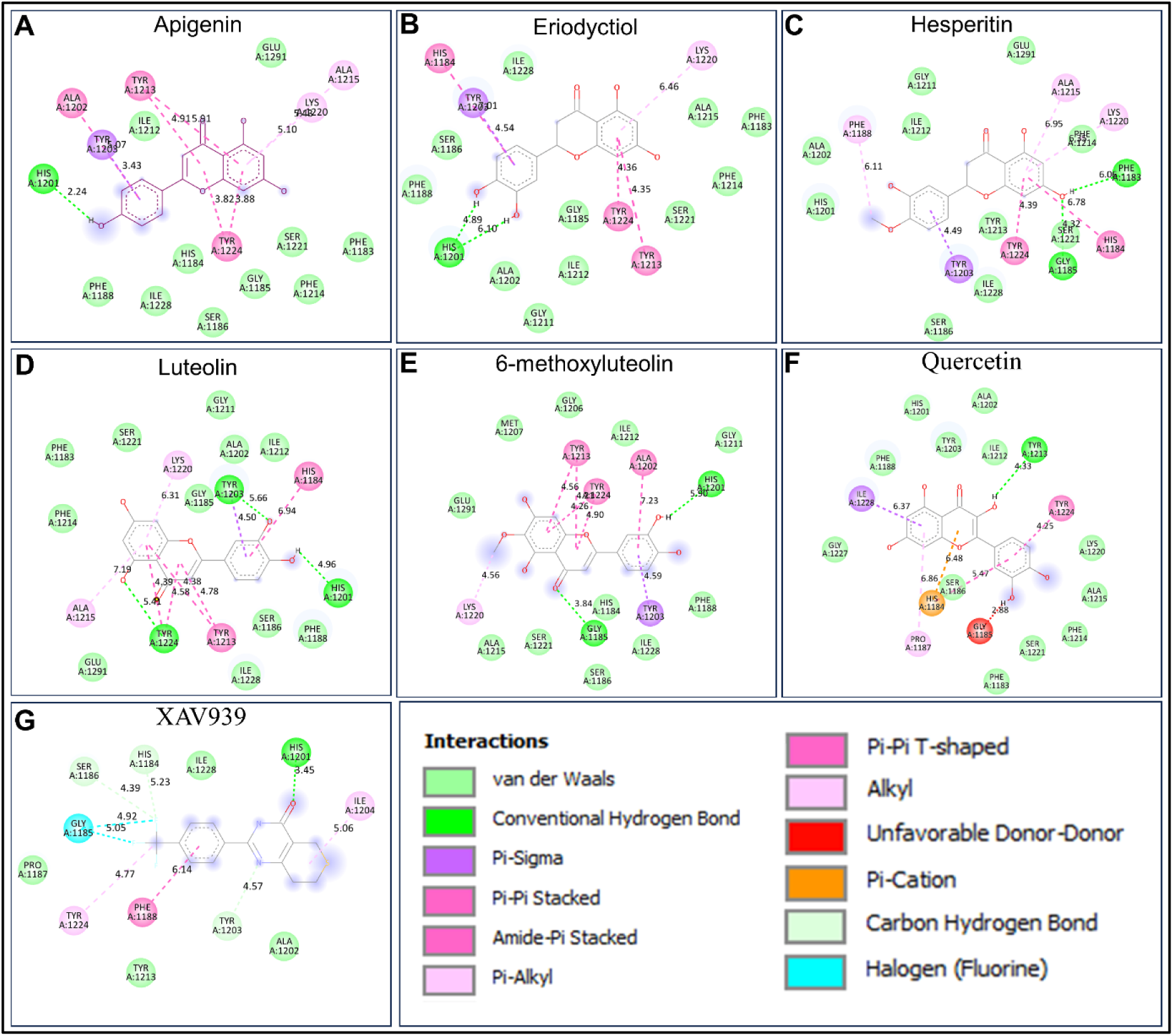
Molecular docking interactions (2-dimensional representations) revealed ligand-protein interactions between tankyrase 1 and olive polyphenols. Interaction of tankyrase 1 with olive phytochemicals are demonstrated (A-F). Tankyrase with Apigenin (A), Eriodyctiol (B), Hesperetin (C), Luteolin (D), 6-methoxyluteolin (E), Quercetin (F). Interaction of tankyrase 1 with XAV939, as reference compound is depicted in G.

**Table 3a:**
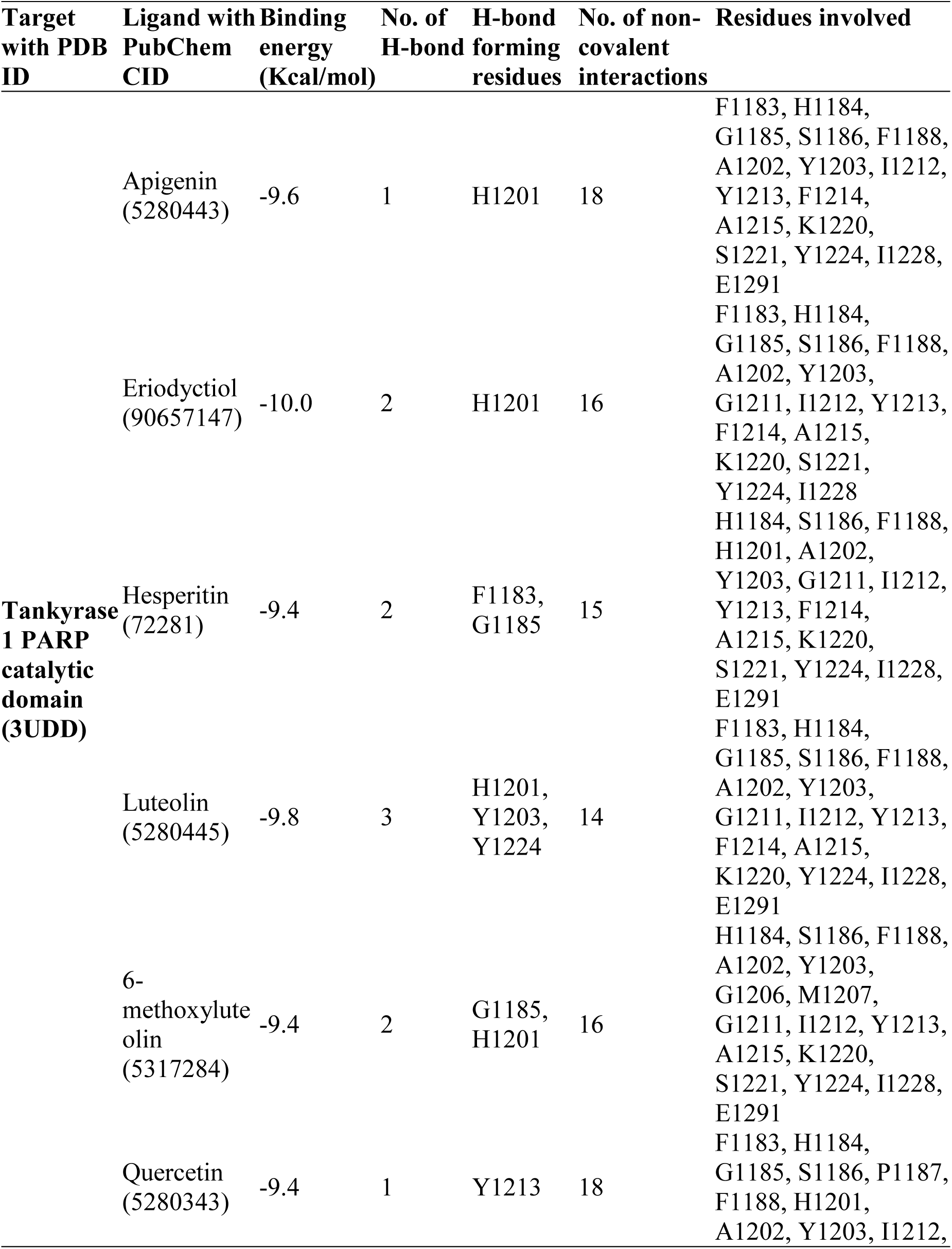
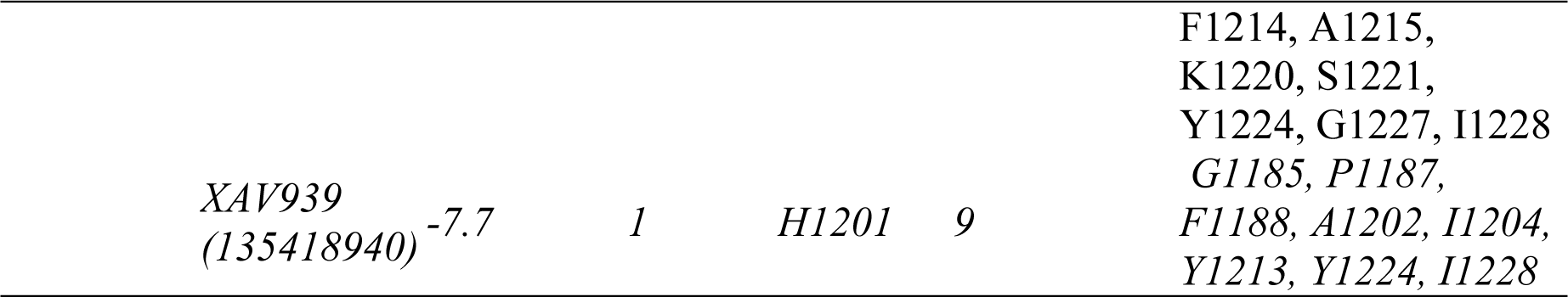
Molecular docking analysis of olive derived compounds and tankyrase 1 PARP c atalytic domain.

The binding energy for each phenolic compound from docking studies surpassed that of the reference compound, XAV939, indicating stronger binding affinity towards tankyrase 1 (Table 3a).

The top 6 compounds selected from the docking study were further docked with the ankyrin repeat cluster domain (TERF1 interacting domain) (PDB ID: 6URQ) of the tankyrase 1 enzyme. Six compounds interacted with the tankyrase 1 ankyrin repeat cluster (ARC) domain by hydrogen bonds, hydrophobic bonds, and Van der Walls interactions. ARC2-3, the second and third ankyrin-repeat clusters of the tankyrase 1 interacts with the TRF1 (telomere repeat factor 1) to induce PARylation of the TRF1 to release it from the telomere which in turn helps telomerase to get access to telomere and maintain telomere stability and continued cell cycle. Thus, blocking the tankyrase activity may help to disrupt the cell cycle in tumor cells. The binding free energy and interactions of target compounds with tankyrase1 ankyrin repeat clusters (ARC2-3) are shown in Table 3b. Apigenin exhibited a good binding affinity (-7.1 Kcal/mol). Apigenin is further involved in Pi-Pi stacked, Pi-Alkyl, carbon hydrogen bonds, and Van der Walls interactions with several residues (Table 3b and Supplementary Fig. 2A). Similarly, other compounds, including eriodyctiol, luteolin, 6-methoxyluteolin, hesperitin, and quercetin also showed good binding affinity by forming H-bonds and non-bonded interactions (Table 3b and Supplementary figure 2). Therefore, these compounds might interact with the ARC2-3 domain of the tankyrase 1 enzyme to disrupt the tankyrase 1-TRF1 interaction, adding another dimension to their anti-cancer mechanisms.

**Table 3b:** Molecular docking analysis of olive derived compounds and tankyrase 1 ankyrin-repeat clusters (ARC2-3).

| Target with<br>PDB ID | Ligand with<br>PubChem<br>CID | Binding<br>energy<br>(Kcal/mol) | No. of H-bond | H-bond<br>forming<br>residues | No. of non-covalent<br>interactions | Residues<br>involved |
| --- | --- | --- | --- | --- | --- | --- |
| **Tankyrase 1**<br>**ankyrin-**<br>**repeat**<br>**clusters**<br>**(ARC2-3)**<br>**(6URQ)** | Apigenin<br>(5280443) | -7.1 | 2 | N408 | 10 | R344, R368,<br>S370, H374,<br>L375, Y379,<br>Y412 |
|  | Eriodyctiol<br>(90657147) | -6.7 | 5 | S370,<br>Y379,<br>N408,<br>Y412 | 8 | R368, L403,<br>H374, L375,<br>D399, S411,<br>E441, K445 |
|  | Hesperitin<br>(72281) | -6.5 | 3 | S370,<br>N408,<br>Y412 | 9 | R368, H374,<br>L375, Y379,<br>D399, L403,<br>S411, E441,<br>K445 |
|  | Luteolin<br>(5280445) | -6.6 | 3 | L455,<br>K504 | 8 | S456, H457,<br>F488, H491,<br>S492, Q495,<br>R498 |
|  | 6-methoxyluteolin<br>(5317284) | -6.6 | 2 | L455,<br>K504 | 9 | S456, H457,<br>F488, H491,<br>S492, Q495,<br>R498 |
|  | Quercetin<br>(5280343) | -6.5 | 4 | D337,<br>R344,<br>D366,<br>R368 | 11 | L340, E341,<br>R344, A364,<br>S365, G367,<br>R368, K369,<br>S370, L375 |

### 3.5. Validation of protein-ligand interactions by Molecular dynamics simulation (MDS) analysis

For further validation of virtual molecular docking results of the PARP catalytic domain and ligands, olive phenols (top six compounds) and known inhibitor XAV939 were subjected to 100 ns MDS. From the MDS trajectory, various parameters were analyzed including the root mean square deviation (RMSD), root mean square fluctuation (RMSF), paired distance between protein and compounds, the radius of gyration (Rg), number of hydrogen bonds, and secondary structure to verify the receptor-ligand conformation properties such as stability and flexibility.. The RMSD values of all six complexes and the reference compound XAV939 (Figure 6A) demonstrated that the selected compounds form stable complexes throughout the simulation, as suggested by better RMSD values compared to XAV939 (Table 4). The overall average RMSF values (Table 4) of receptor amino acid residues for all the compounds from the MDS study further revealed stable interaction with the receptor protein. Additionally, the strength of inhibitors binding to tankyrase 1 was investigated through pair distance analysis. The distance between tankyrase 1 and potential inhibitors was extracted from the MD trajectory (Figure 6C). The critical distance maintained by the inhibitors with target protein tankyrase 1 was ∼ 0.2 nm (2 Å), which indicates a robust short-range interaction between protein and olive phenolic compounds. Rg, which is an indicator of compactness, stability, and folding of structure, was calculated based on the intrinsic dynamics of protein-ligand complexes. All the complexes had a steady average Rg value of ∼1.72 nm over the 100 ns MD simulation. Next, we recorded the hydrogen bonding dynamics throughout the 100 ns simulations, as it is one of the most critical non-covalent interactions. The number of hydrogen bonds is directly correlates to the interaction strength. The average number of hydrogen bonds was between 0.7 and 1.7, where most hydrogen bonds formed in the case of tankyrase 1-quercetin complex.

**Figure 6:**
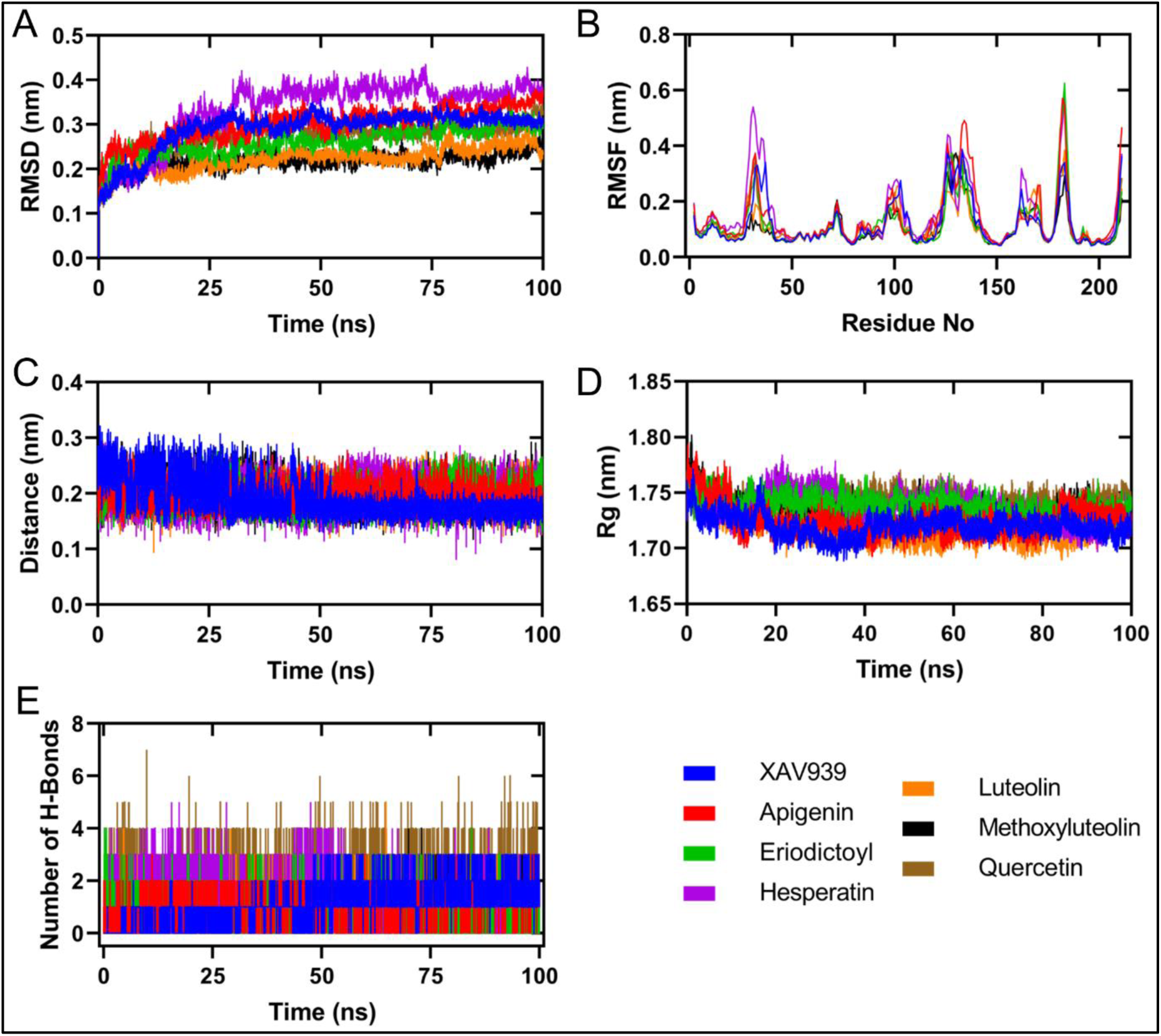
MD simulation parameters of the study between tankyrase 1 and olive-derived compounds. (A) Root means square deviation (RMSD) of tankyrase 1 upon binding of olive phytochemicals over 100 ns. (B) Root means square fluctuation (RMSF) of tankyrase 1 in residual level over 100 ns. (C) Pairing distance between tankyrase 1 and inhibitor molecules depicted over 100 ns simulation. (D) Radiation of gyration (Rg) of tankyrase 1 upon binding of inhibitor over 100 ns. (E) Number of hydrogen bonds throughout 100 ns simulation.

**Table 4:** Average of MD parameters including standard deviations of TNKS1-inhibitor complexes.

|  |  | **XAV9**<br>**39** | **Apigen**<br>**in** | **Eriodyct**<br>**iol** | **Hespere**<br>**tin** | **Luteol**<br>**in** | **6-**<br>**methoxylute**<br>**olin** | **Querce**<br>**tin** |
| --- | --- | --- | --- | --- | --- | --- | --- | --- |
| **RMS**<br>**D**<br>**(nm)** | Mea<br>n | 0.2891 | 0.3015 | 0.2631 | 0.3347 | 0.2218 | 0.2176 | 0.2736 |
| **RMS**<br>**F**<br>**(nm)** | Mea<br>n | 0.1209 | 0.1398 | 0.1181 | 0.1495 | 0.1111 | 0.1029 | 0.1188 |
| **Paire**<br>**d**<br>**distan**<br>**ce**<br>**(nm)** | Mea<br>n | 0.1893 | 0.1913 | 0.1932 | 0.2042 | 0.2074 | 0.1922 | 0.185 |
| **Rg**<br>**(nm)** | Mea<br>n | 1.72 | 1.725 | 1.739 | 1.737 | 1.716 | 1.732 | 1.743 |
| **H-**<br>**bond** | Mea<br>n | 1.23 | 1.005 | 0.9753 | 1.176 | 0.6987 | 0.7734 | 1.773 |

Further, we also checked the binding conformation of selected ligands within the binding pocket of tankyrase 1 by taking a snapshot of each 20 ns (Figure 7), and all the ligands were found to be bound within the binding pocket for the entire simulation period (100 ns).

**Figure 7:**
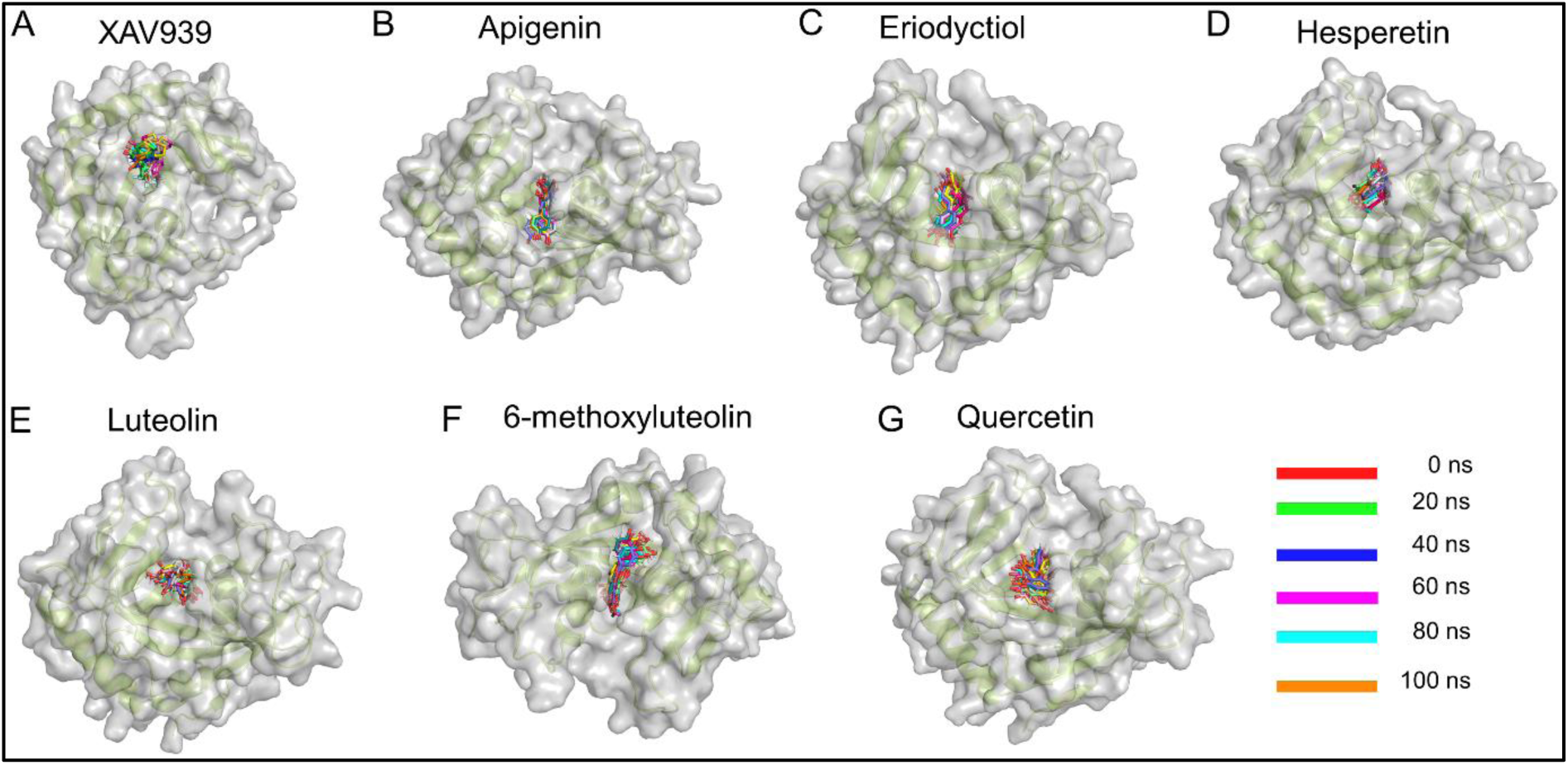
Binding conformation of inhibitors in tankyrase 1 with 20 ns time interval. The olive tree-derived compounds are depicted in different colors inside the binding pocket based on the time frame: red, 0^th^ ns; green, 20^th^ ns; blue, 40^th^ ns; magenta, 60^th^ ns; cyan, 80^th^ ns; orange, 100^th^ ns.

### 3.6. Binding free energy analysis

In addition, the binding energy of receptor-ligand complexes was also calculated using MMPBSA methods throughout the MD simulation. The g_mmpbsa module is employed to calculate the binding free energy, which includes Van der Waal’s, electrostatic interaction, and polar solvation energies. Van der Waal’s and electrostatic interactions are two of the key players in receptor-drug interactions. Electrostatic interactions pull the drug towards the binding pocket, and van der Waals interaction keeps the drug inside the binding pocket. Most of the selected drugs (apigenin, eriodyctiol, luteolin, 6-methoxyluteolin, and quercetin) showed better binding energy, whereas hesperetin showed similar binding energy with the reference compound XAV939 (Table 5). Thus, all the compounds can be considered better inhibitors than XAV939. Overall, these compounds might be in the best position among olive polyphenols to interfere in tankyrase 1 enzymatic action and subsequent repression of the Wnt/β-catenin pathway. However, we selected only the three top compounds based on binding free energy (Table 5) (apigenin, quercetin, and luteolin) for further *in vitro* studies.

**Table 5:**
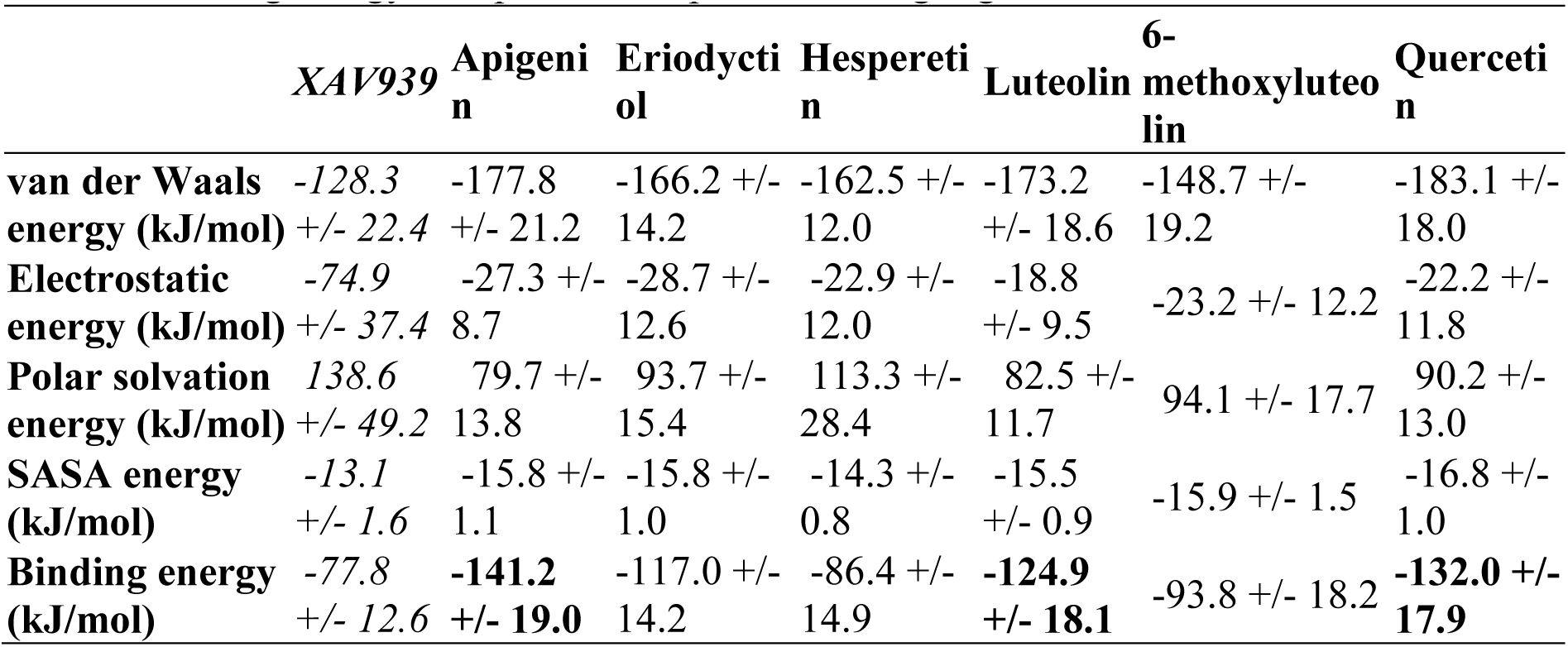
Binding free energy analysis of Tankyrase 1-inhibitor complexes by MM-PBSA method. Binding energy of top three compounds are highlighted in bold.

|  | *XAV939* | **Apigenin** | **Eriodyctol** | **Hesperetin** | **Luteolin** | **6-methoxyluteolin** | **Quercetin** |
| --- | --- | --- | --- | --- | --- | --- | --- |
| **van der Waals energy (kJ/mol)** | -128.3 +/- 22.4 | -177.8 +/- 21.2 | -166.2 +/- 14.2 | -162.5 +/- 12.0 | -173.2 +/- 18.6 | -148.7 +/- 19.2 | -183.1 +/- 18.0 |
| **Electrostatic energy (kJ/mol)** | -74.9 +/- 37.4 | -27.3 +/- 8.7 | -28.7 +/- 12.6 | -22.9 +/- 12.0 | -18.8 +/- 9.5 | -23.2 +/- 12.2 | -22.2 +/- 11.8 |
| **Polar solvation energy (kJ/mol)** | 138.6 +/- 49.2 | 79.7 +/- 13.8 | 93.7 +/- 15.4 | 113.3 +/- 28.4 | 82.5 +/- 11.7 | 94.1 +/- 17.7 | 90.2 +/- 13.0 |
| **SASA energy (kJ/mol)** | -13.1 +/- 1.6 | -15.8 +/- 1.1 | -15.8 +/- 1.0 | -14.3 +/- 0.8 | -15.5 +/- 0.9 | -15.9 +/- 1.5 | -16.8 +/- 1.0 |
| **Binding energy (kJ/mol)** | -77.8 +/- 12.6 | **-141.2 +/- 19.0** | -117.0 +/- 14.2 | -86.4 +/- 14.9 | **-124.9 +/- 18.1** | -93.8 +/- 18.2 | **-132.0 +/- 17.9** |

### 3.7. Olive polyphenols induced cytotoxicity in HT-29 cells

The three top ligands (apigenin, luteolin, and quercetin) were further studied in an *in vitro* to investigate the anticancer effects of these compounds. First, the ability of these three compounds to cause cytotoxicity was studied in HT-29 CC cells. These three compounds reduced HT-29 cell viability in a dose-dependent manner in both 24 h and 48 h of exposure. Apigenin significantly reduced cell viability up to 42% and 46% for the 24 h and 48 h exposures (Figure 8A), respectively, whereas quercetin reduced cell viability up to 42% and 49% for the 24 h and 48 h exposures, respectively (Figure 8B). On the other hand, luteolin significantly reduced cell viability up to 51% and 54% for the 24 h and 48 h exposures, respectively (Figure 8C). We wanted to check this, particularly to get an insight into the combinatorial effects of these compounds (doses were selected that induced ∼50% cell death), if any. In the case of combinatorial treatment, cell death was significantly decreased compared to the individual treatment groups except the luteolin vs luteolin + apigenin group (Figure 8D).

**Figure 8:**
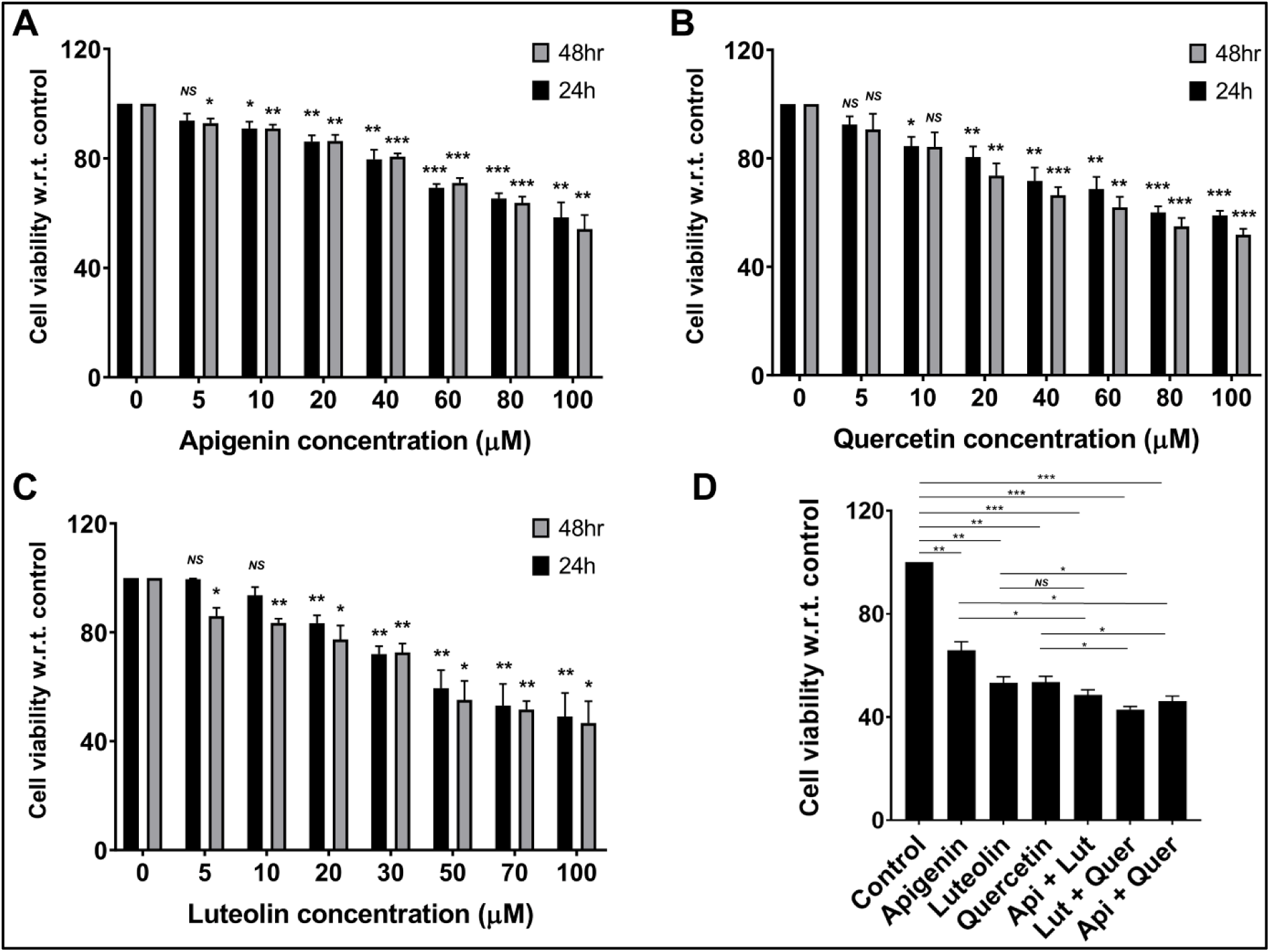
Effects of apigenin, quercetin, and luteolin on HT-29 cell viability. Cell viability was reduced significantly upon treatment with apigenin (A), quercetin (B), luteolin (C) alone, for 24 h and 48 h or in combination for 48 h (D) in a dose dependent way. In the case of combination treatment apigenin and quercetin was used at 80 µM concentration, whereas luteolin was used at 70 µM concentration (48 h exposure time). (\**P* < 0.05, \*\**P*< 0.01, \*\*\**P*< 0.001 vs untreated group, data represented as the mean ± SEM of at least three independent experiments.

### 3.8. Olive polyphenols reduced colony formation ability of the HT-29 cells

Next, we studied the colony-forming ability of CC cells in the presence of the top three selected compounds. Quercetin and luteolin reduced the number of colonies after 48 hours of exposure. Additionally, selected polyphenols reduced the colony number more when used together than the single compound treatment (Figure 9). As olive oil showed preventive action against CC, these compounds might act in combination, and we wanted to check for combinatorial activities of these compounds as well. Our results indicated that the selected polyphenol compounds induced cell death and prevented single cells from remaining attached to the surface and growing into individual colonies over time and when used in combination induced increased cytotoxic activities (Figure 9).

**Figure 9:**
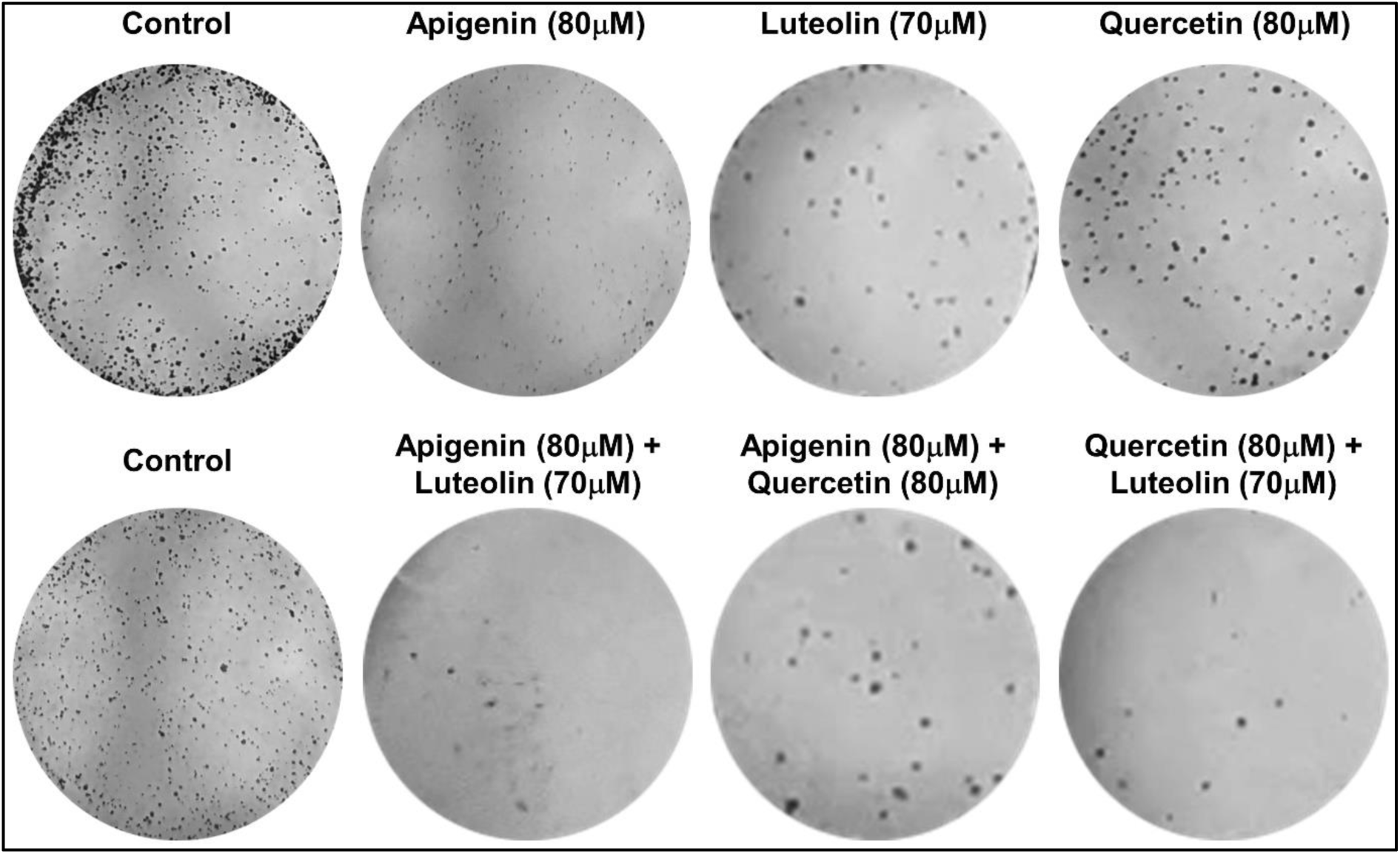
Olive polyphenols either alone or in combination reduces colony formation of HT-29 cells. Representative image of the effect of apigenin, quercetin, luteolin, and the combination of these compounds on colony formation ability of the HT-29 cells. HT-29 cells were treated for 48 h either alone or in combination and then allowed to grow into colonies. After 10 days’ cells were fixed and stained with crystal violet and a significant reduction in number of colonies was observed in case of treatment groups.

### 3.9. Olive polyphenols induced apoptosis in HT-29 cells

Next, we aimed to unravel the mechanism by which the top selected compounds cause cytotoxicity, and we found that the top three selected compounds, apigenin, quercetin, and luteolin, either alone or in combination, induced apoptosis in HT-29 cells when treated for 48 h. AO/EtBr dual staining reveals the apoptosis-related changes of the cell membrane when checked under a fluorescence microscope. Apoptosis was not detected in the control group (without treatment) (Figure 10A). Early-stage apoptotic cells (crescent-shaped or granular yellow-green AO nuclear staining) were detected in apigenin, quercetin, luteolin, or combination groups (Figure 10B - 10G). On the other hand, late-stage apoptotic cells were detected in treatment groups characterized by concentrated and asymmetrically localized orange nuclear EtBr staining (Figure 10B - 10G). The number of necrotic cells was also increased in volume, with uneven orange-red fluorescence at their periphery (Figure 10). In combination groups, cells appeared to be in the process of disintegrating.

**Figure 10:**
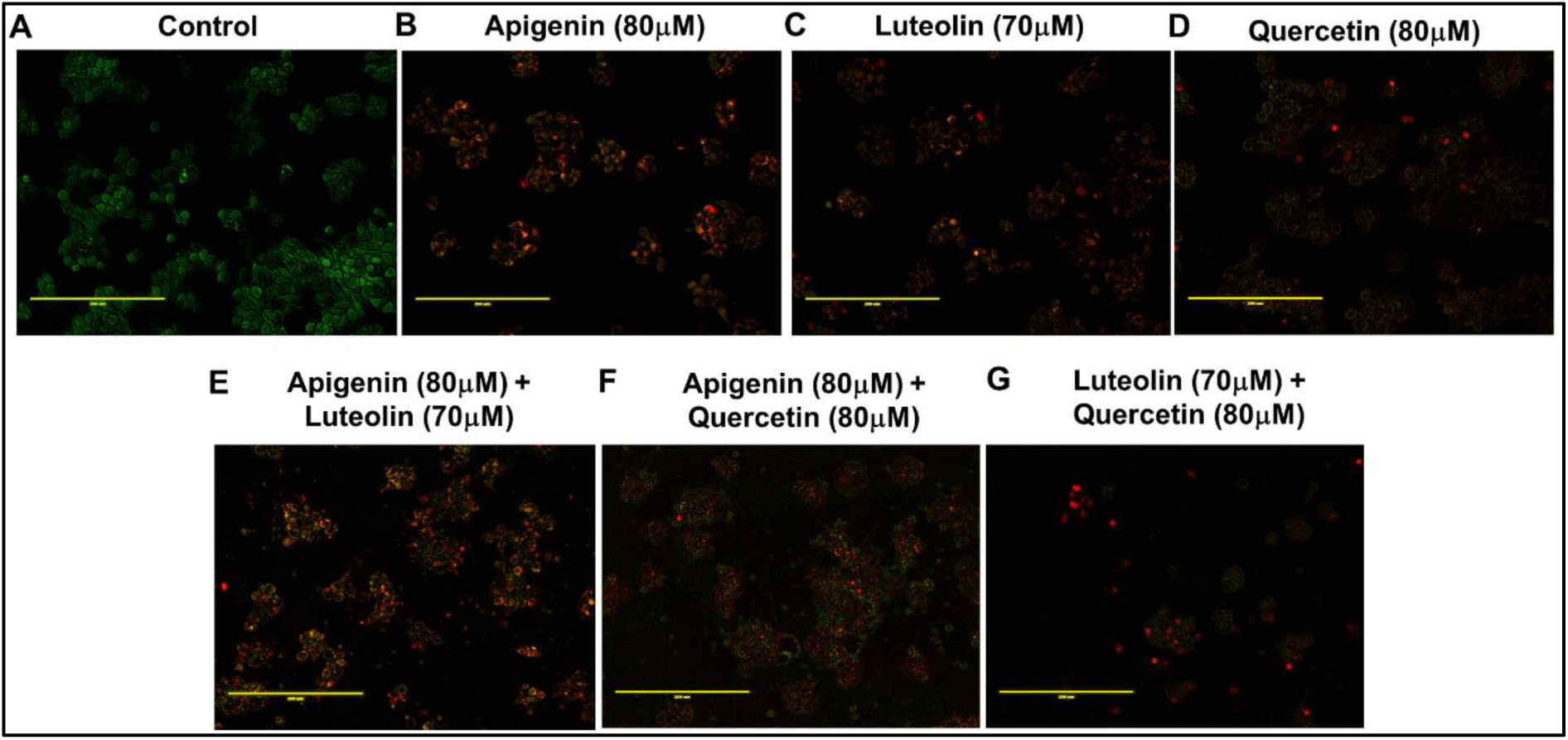
Selected olive polyphenols induced apoptosis and necrosis in HT-29 cells. HT-29 cells either left untreated (A) or treated with alone with apigenin (B), luteolin (B) quercetin (C) or in combination (E, F and G) for 48 h and then AO/EtBr solution was added to assess apoptosis. Green stain indicates the healthy live cells whereas red stain indicates apoptotic, necrotic, and dead cells. Fluorescence microscopic pictures were taken at 200X magnification.

### 3.10. Olive polyphenols suppress the β-catenin downstream gene expression

Further, we wanted to check the effect of tankyrase inhibition by apigenin, luteolin, and quercetin on Wnt/β-catenin pathway and especially the expression of genes regulated by β-catenin accumulation in cytoplasm and subsequent translocation into nucleus. The mRNA level expression of few of the most important genes regulated by β-catenin were evaluated including cyclin D1 (*CCND1*), proto-oncogene c-Myc (*MYC*), and cyclin-dependent kinase 4 (*CDK4*). Tankyrase 1 inhibition will suppress the β-catenin nuclear translocation, which will in turn block downstream gene expression. Here, we found selected compounds significantly reduced the total protein level of β-catenin in HT-29 cells (11A), and gene expression of *CCND1* (Figure 11B), *MYC* (Figure 11C), and *CDK4* (Figure 11D) also diminished in treated cells compared to the untreated cells, either alone or in combination. These results indicate that the Wnt/β-catenin axis gets inhibited by these compounds by blocking the tankyrase 1 activity. Apigenin and luteolin reduced *CCND1* expression by ∼30% (Figure 11B), whereas quercetin reduced *CCND1* expression by ∼70% (Figure 11B) in treated HT-29 cells. In the case of combination treatment, *CCND1* expression was further reduced in treated cells, especially in the combination of apigenin-quercetin and luteolin-quercetin treatments, ∼ 80% and 90%, respectively (Figure 11B). Similarly, apigenin, luteolin, and quercetin decreased *MYC* expression by around 25%, 35%, and 55% (Figure 11C), respectively, in treated cells compared to untreated cells, whereas, the combination of apigenin-luteolin, apigenin-quercetin, and luteolin-quercetin reduced *MYC* expression in treated cells by around 70%, 80%, and 90%, respectively (Figure 11C). These compounds also significantly suppressed *CDK4* expression. Treatment of HT-29 cells with olive polyphenols apigenin, luteolin, and quercetin reduced *CDK4* expression up to 35%, 25%, and 40%, respectively (Figure 11D).

**Figure 11:**
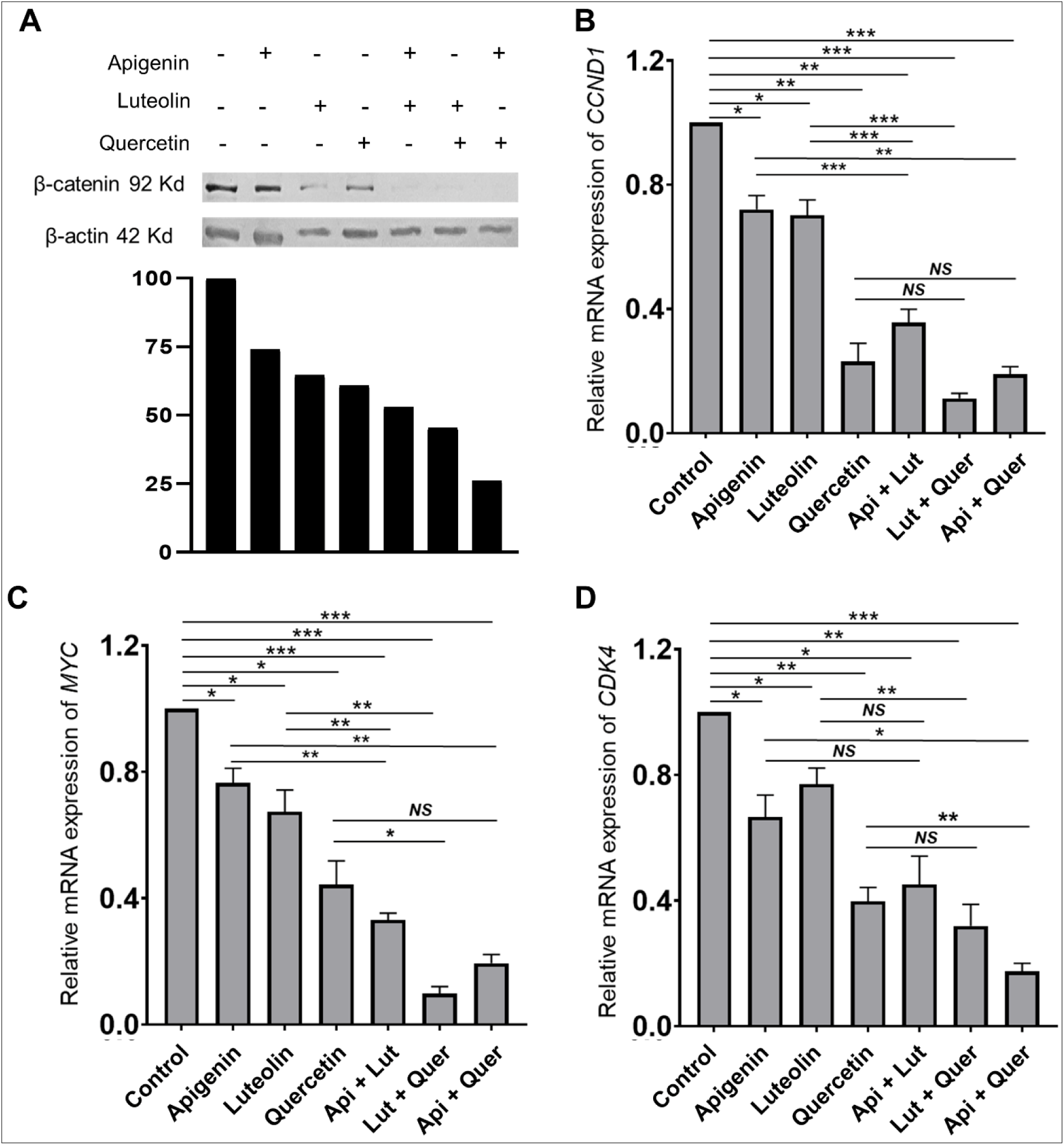
Selected olive polyphenols reduced total β-catenin and also suppressed downstream gene expression of tankyrase 1/Wnt/β-catenin axis: HT-29 cells were treated with selected phytochemicals either alone (apigenin 80 µM/quercetin 80µM/ and luteolin 70 µM) or in combination (Apigenin + Luteolin, Luteolin+ Quercitin, Apigenin+ Quercitnin) for 48 h. A) Upper panel demonstrated the total β-catenin protein level, checked by Immunoblot analysis. β-actin was employed as housekeeping protein. Lower panel depicted the densitometry analysis by Image J where band density of both β-catenin and β –actin protein bands from three independent experiments were considered and average ratio of β-catenin: β –actin intensity was plotted. B-D) RT-PCR analysis to demonstrate the relative gene expression of cyclin D1 (CCND1) (B), proto-oncogene c-Myc (MYC) (C), and cyclin-dependent kinase 4 (CDK4) (D) in above mentioned treatment. These are representation of three independent experiments. (\**P* < 0.05, \*\**P*< 0.01, \*\*\**P*< 0.001 vs untreated group, data represented as the mean ± SEM of at least three independent experiments.

Further, a reduction in *CDK4* expression was also observed when these compounds were treated in combination. The combination of apigenin-luteolin, apigenin-quercetin, and luteolin-quercetin reduced *CDK4* expression by almost 55%, 70%, and 85% in treated cells compared to untreated cells (Figure 11D). Overall, these compounds successfully suppressed the growth-promoting activities of Wnt/β-catenin by blocking tankyrase 1 action. Moreover, when these compounds are present in a combination, such as enriched sources like olive, it might be more beneficial against the CC.

### 3.11. Olive polyphenols restrained CRC spheroid formation

In a 3D culture system, the selected compounds apigenin, luteolin, and quercetin demonstrated significant effectiveness in restraining HT-29 spheroid formation. When these compounds were introduced into the culture media, either individually or in combination, in ultra-low attachment plates, a notable reduction in spheroid growth, both in size and cell viability was observed (Figure 12). The extent of their effectiveness was found to vary based on the duration of exposure. In the case of combination treatment, a more pronounced inhibitory effect was observed. This suggests that when these compounds are present together in a single food source, their beneficial effects might be enhanced, indicating the potential development of combination therapies. The synergistic impact of the compounds in a combined treatment approach could offer enhanced therapeutic outcomes in the context of colon cancer. These results highlight the potential anti-cancer properties of apigenin, luteolin, and quercetin against HT-29 cells in a three-dimensional cellular environment.

**Figure 12:**
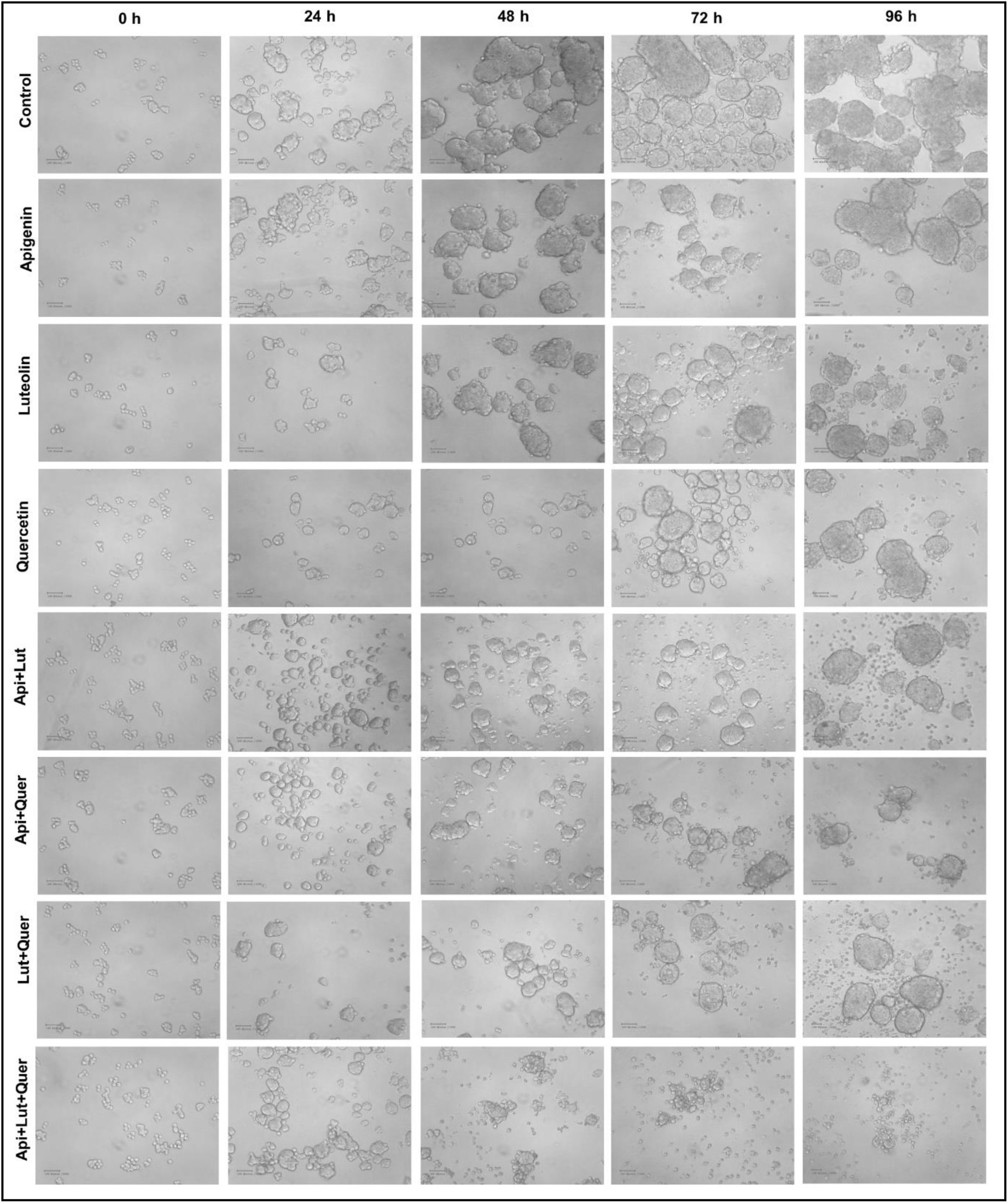
Apigenin, luteolin, and quercetin inhibited HT-29 spheroid formation. HT-29 cells were seeded at very low concentration in 12-well ultra-low attachment plate in serum free DMEM/F12 stem cell media. Selected olive phytochemicals either alone (apigenin 80 µM, quercetin 80µM, and luteolin 70 µM) or in combination were administered after overnight incubation and spheroids were allowed to grow. Spheroid morphology and growth were observed under inverted microscope and images were captured (100X) every 24 h gap. All the phytocompounds reduce the number of spheroids as well as the size of individual spheroid, however combinational treatment found to be more effective than individual treatment in inhibiting spheroid formation, whereas simultaneous treatment with all three compounds at the same time exhibited best effect in spheroid size and number reduction.

Overall, this study further characterized tankyrase 1 as an important target in colon cancer. On top of that, we have identified the potential anti-colon cancer mechanism of olive polyphenols in the form of targeting and inhibiting tankyrase 1. Three phenolic compounds, including apigenin, luteolin, and quercetin, were identified as the best hits to target tankyrase 1 among the olive phytochemicals, and possibly these compounds are responsible for the anti-colon cancer activities of olive.

## Author contributions

AS and DN conceived the idea and designed the experimental strategy. AS, SB, DK, MS and MV performed the *in silico* experiment. AS performed the *in vitro* experiment. PS and BS assisted in the *in vitro* experimentation. the study protocol. AS, SB and DN interpreted the results. AS, SB and DK prepared the figures and tables. AS prepared the manuscript while SB and DK assisted the Molecular Docking and Molecular Dynamics simulation study part. DN corrected and revised the manuscript at its final version.

All authors critically read, and approved the final manuscript.

## Acknowledgment

AS Acknowledge the support from Council of Scientific and Industrial Research (CSIR), Govt. of India for fellowship for doctoral research. PS, BS, MS and MV were supported by fellowship from Department of Biotechnology, Govt. of India. The authors acknowledge the help of Prof. Jaya Bandyopadhyay and her lab members for providing fluorescence microscope facility.

## Funding

This work was supported by Department of Science and Technology, Govt. of India (DST)-Inspire faculty research grant (DST/INSPIRE/04/2017/000675) and Maulana Abul Kalam Azad University of Technology, West Bengal (MAKAUT, WB) research seed grant to DN.

## Data Availability Statement

All data generated or analyzed during this study are included in this published article.

## Disclosure Statement

Authors declares no competing interest.

## Notes

### Competing Interest Statement

The authors have declared no competing interest.

