## Supplementary figure 1 for "Olive phenolic compounds, potent tankyrase 1 inhibitor exhibit anti colon cancer effects by blocking Wnt/β-catenin pathway"

### Slide 1
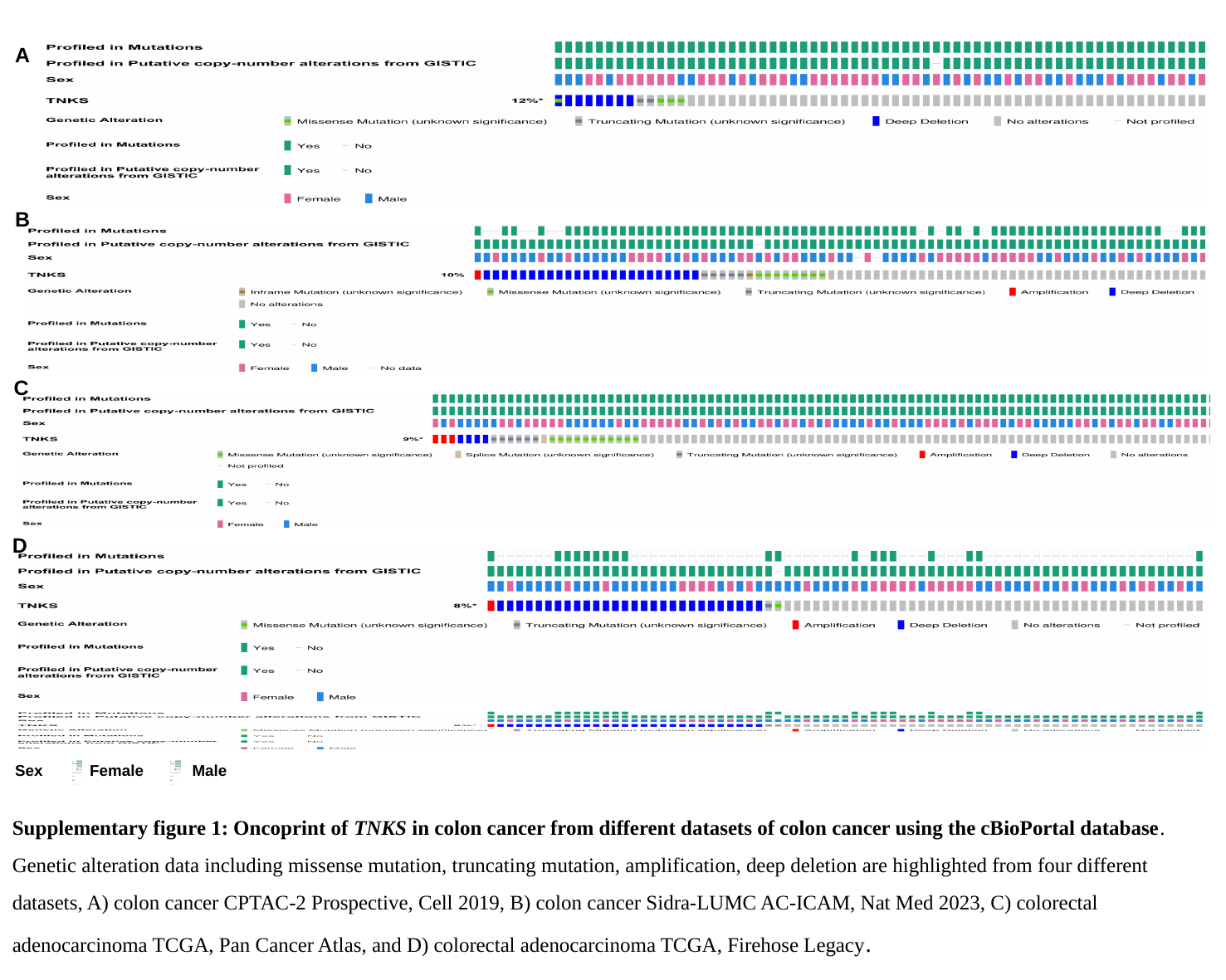

A
B
C
D
Sex
Female
Male
Supplementary figure 1: Oncoprint of TNKS in colon cancer from different datasets of colon cancer using the cBioPortal database. Genetic alteration data including missense mutation, truncating mutation, amplification, deep deletion are highlighted from four different datasets, A) colon cancer CPTAC-2 Prospective, Cell 2019, B) colon cancer Sidra-LUMC AC-ICAM, Nat Med 2023, C) colorectal adenocarcinoma TCGA, Pan Cancer Atlas, and D) colorectal adenocarcinoma TCGA, Firehose Legacy.
