## Supplementary figure 2 for "Olive phenolic compounds, potent tankyrase 1 inhibitor exhibit anti colon cancer effects by blocking Wnt/β-catenin pathway"

### Slide 1
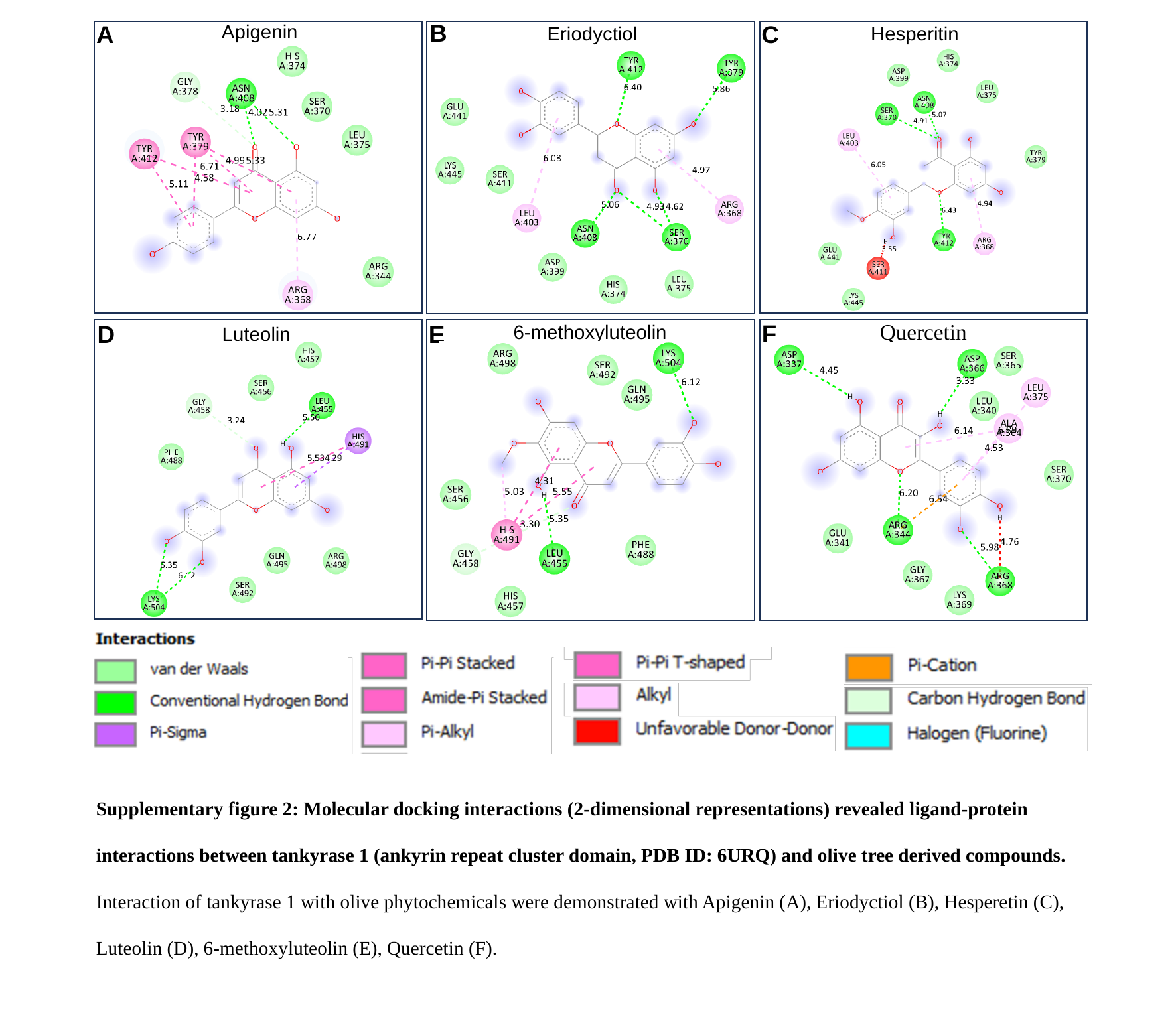

B
C
A
Apigenin
Eriodyctiol
Hesperitin
Quercetin
F
E
D
6-methoxyluteolin
Luteolin
Supplementary figure 2: Molecular docking interactions (2-dimensional representations) revealed ligand-protein interactions between tankyrase 1 (ankyrin repeat cluster domain, PDB ID: 6URQ) and olive tree derived compounds. Interaction of tankyrase 1 with olive phytochemicals were demonstrated with Apigenin (A), Eriodyctiol (B), Hesperetin (C), Luteolin (D), 6-methoxyluteolin (E), Quercetin (F).
