## Supplementary table 2 for "Olive phenolic compounds, potent tankyrase 1 inhibitor exhibit anti colon cancer effects by blocking Wnt/β-catenin pathway"

**Supplementary Table 2:** Binding free energy of the tankyrase 1 and olive phytochemicals. Reference compound is shown in italics.

| **Protein** | **Ligand** | **Binding affinity (Kcal/mol)** |
| --- | --- | --- |
| **Tankyrase 1**  **[3UDD]** | Apigenin | -9.6 |
|  | Apigenin-7-o-glucoside | -8.9 |
|  | Eriodictyol | -10.0 |
|  | Hesperitin | -9.4 |
|  | Luteolin | -9.8 |
|  | 6-methoxyluteolin | -9.4 |
|  | Quercetin | -9.4 |
|  | Berchemol | *-6.8* |
|  | Chlorogenic Acid | *-7.8* |
|  | 3,4-Dihydroxyphenylglycol | *-6.7* |
|  | Loganic Acid | *-6.7* |
|  | Loganin | *-7.8* |
|  | Taxifolin | *-9.0* |
|  | Gentisic Acid | *-6.8* |
|  | Cornoside | *-7.3* |
|  | Chrysoeriol | *-9.1* |
|  | Rosmarinic Acid | *-9.0* |
|  | *XAV939* | *-7.7* |
